# Comparative transcriptomics of *Lathyrus sativus* reveals accession-specific resistance responses against *Erysiphe pisi*

**DOI:** 10.1101/2024.11.23.624375

**Authors:** Rita Maravilha Marques, Telma Fernandes, Pedro M. Barros, Susana Trindade Leitão, Diego Rubiales, Maria Carlota Vaz Patto, Carmen Santos

## Abstract

*Lathyrus sativus* (grass pea) is a valuable crop for sustainable agriculture, offering both dietary benefits and desirable agronomic traits. However, its yield stability is limited by different diseases such as powdery mildew caused by *Erysiphe pisi*. Frequent fungal resistance to pesticides and growing environmental concerns highlight the need for research investment to develop resistant crop varieties. Four *L. sativus* accessions, exhibiting varying levels of resistance to *E. pisi* (resistant, partially resistant, partially susceptible, and susceptible), were analysed using dual RNA-seq to identify key defence mechanisms and effector genes involved in this plant-pathogen interaction. The dual transcriptomic analysis highlighted a host biphasic response, characterised by an initial burst of gene expression, followed by a quiescent phase, and a second wave of intense gene expression at 72 hours after inoculation. Common *L. sativus* defence mechanisms, including antifungal protein expression, cell wall reinforcement, reactive oxygen species-mediated defence were activated by all accessions compared to susceptible accession. Unique responses in the resistant accession integrate early reinforcement of structural barriers with sustained chemical defences and stress responses. Overall, the partially resistant accession exhibited a front-loaded defence response, focused on biotic stimuli and interspecies interactions at early infection stages. In contrast, partial susceptible accessions exhibited a weaker baseline defence system, with a slower and less robust response specifically targeting pathogen infection. We identified potential *E. pisi* effectors, including genes involved in cell wall hydrolysis, nutrient acquisition, and virulence, with a higher diversity of effectors identified in the susceptible accession. This study identifies novel targets within the complex defence mechanisms of the *Lathyrus sativus*-*Erysiphe pisi* interaction that will support to future breeding programs aimed at enhancing resistance to *E. pisi* in *L. sativus* and other related species.

## 1. Introduction

*Lathyrus sativus* (grass pea) is an annual legume highly valued both as a nutritious food source for humans and as animal feed. As the most widely cultivated species within the *Lathyrus* genus, it combines significant dietary benefits with desirable agronomic traits (Peña-Chocarro & Peña, 1999; Lambein et al., 2019). *Lathyrus sativus* cultivation demands minimal inputs, demonstrating remarkable resilience, in challenging conditions such as drought, flooding, and poor soils (Rubiales et al., 2020; Gonçalves et al., 2022; Sanches et al., 2024). It is traditionally cultivated in drought-prone, marginal areas of South Asia and East Africa (Vaz Patto & Rubiales, 2014). In Mediterranean regions, *L. sativus* is vital in supporting local economies (Gonçalves et al., 2015; Rubiales et al., 2020). Certain *L. sativus* accessions exhibit high resistance to air and soil-born fungal diseases (Vaz Patto et al., 2006; Vaz Patto & Rubiales, 2009, 2014; Sampaio et al., 2021; Martins et al., 2022; Martins et al., 2023), making them valuable sources of resistance genes. Due to their phylogenetic proximity, *Lathyrus* spp. share many pathogens with *Pisum*, *Lens*, and *Vicia* genera that include crops like pea, lentil, and vetches.

Powdery mildew is one of the most widespread and damaging airborne fungal diseases (Sulima & Zhukov, 2022). In legumes, it is caused by obligate biotrophic ascomycetes from the order Erysiphales (Rubiales et al., 2015; Martins et al., 2020; Sulima & Zhukov, 2022). Powdery mildew on *Lathyrus* spp. is often caused by *Erysiphe pisi* (Vaz Patto et al., 2006; Martins et al., 2020), the same pathogen responsible for the disease in pea (*P. sativum*), which also infects species within the *Medicago*, *Vicia*, *Lupinus,* and *Lens* genera (Fondevilla & Rubiales, 2012). The management of powdery mildew in the field has traditionally relied on chemical fungicides. However, increasing resistance among fungal strains and increasing environmental concerns have pushed for alternative control strategies, including the breeding of disease-resistant varieties (Fondevilla et al., 2007; Mundt, 2014).

Plant disease resistance has been divided broadly in two categories: incomplete or partial resistance provided by quantitative disease resistance (QDR) genes and complete resistance mediated by resistance genes (R-genes) (Delplace et al., 2022). R gene resistance mediates the plant immune system, which has two layers. The first relies on pattern recognition receptors (PRRs) that detect pathogen-associated molecular patterns (PAMPs), activating PAMP-triggered immunity (PTI) (Zipfel & Robatzek, 2010; Roux et al., 2014; Lolle et al., 2020). However, pathogens can bypass this defence by secreting effectors that suppress PTI and hijack host proteins (Vleeshouwers & Oliver, 2014). In response, plants use intracellular resistance receptors such as nucleotide-binding leucine-rich domain proteins (NLRs) to recognise effectors, activating a stronger defence response called effector-triggered immunity (ETI) (Roux et al., 2014; Adachi et al., 2019; Li et al., 2020). This recognition leads to prolonged resistance through various immune response pathways like reactive oxygen species (ROS) production, hypersensitive cell death (HR), and systemic-acquired resistance (SAR) (Adachi et al., 2019; Li et al., 2020b; Derevnina et al., 2021).

Research on powdery mildew resistancés genetic and molecular bases has largely focused on model and important cereal crops plants such as Arabidopsis, barley, and wheat (Yun et al., 2016; Zhang et al., 2016; Kuhn et al., 2017; Kusch & Panstruga, 2017; Hoseinzadeh et al., 2019). In contrast, legume crops resistance mechanisms, including *L. sativus*, remain underexplored, hampering a more efficient and effective development of disease-resistant varieties. Previous studies have identified three key genes linked to *E. pisi* resistance in pea: the recessive genes *er1* (also known as *MLO1*) and *er2*, and the dominant gene *Er3.* The HR plays a major role in pea resistance, which is governed by *er2* and *Er3* genes (Fondevilla and Rubiales, 2012; Barilli et al., 2014). In contrast, *er1* provides from complete to moderate levels of resistance by blocking fungus haustoria formation (Fondevilla et al., 2006; Fondevilla and Rubiales, 2012; Iglesias-García et al., 2015). However, concerns about the durability of these resistance genes, due to pathogen evolution, underscore the need for additional resistance sources (Fondevilla et al., 2013). Partial resistance is a potentially more durable approach than complete resistance, due to the reduced selective pressure imposed on the pathogen (McDonald and Linde, 2002; Niks and Rubiales, 2002). In the model legume *Medicago truncatula*, significant progress has been made in identifying quantitative trait loci (QTLs) for *E. pisi* disease symptoms (mycelium and conidia covering the leaf surface) (Ameline-Torregrosa et al., 2008; Yang et al., 2013). QTLs for resistance have been mapped to chromosomes 4 and 5, corresponding to the loci *Epp1* (on chromosome 4), *Epa1*, and *Epa2* (on chromosome 5), while a dominant resistance gene *MtREP1* (resistance to *Erysiphe pisi* race 1), was also identified and mapped on chromosome 5 (Ameline-Torregrosa et al., 2008; Yang et al., 2013). More recently, a dual RNA-Seq approach has provided deeper insights into the interactions between *M. truncatula* and *E. pisi*, revealing R-gene-mediated resistance against involves transcriptional reprogramming, amplifying PTI, activating the jasmonic acid/ethyne signalling network, and balancing growth-defence. Susceptibility is linked to suppressed defence signalling, reduced cell wall defences, and processes favouring biotrophy. (Gupta et al., 2020). Additionally, sugar transporters were found to mediate basal resistance to powdery mildew in *M. truncatula* (Gupta et al., 2021). In *Lathyrus* spp., a genome-wide association study (GWAS) identified 12 single nucleotide polymorphisms (SNPs) linked to disease severity to *E. pisi* in *L. sativus* mapped across all chromosomes, except in chromosome 1 (Martins et al., 2023). In *L. cicera*, three QTLs mapped on linkage groups I, II and IV were associated with partial resistance to *E. pisi* (Santos et al., 2020). The characterisation of the *MLO1* genes in both species revealed that *Lathyrus* MLO1 proteins belong to Clade V, a group linked with powdery mildew susceptibility in dicots (Santos et al., 2021).

Breeding for sustainable durable resistance in crops includes strategies like pyramiding major resistance genes and quantitative resistance genes. NLR proteins, central to ETI, display significant structural diversity and are typically classified into four classes based on their N-terminal domains: coiled-coil NLR (CC-NLR), Toll/interleukin-1 receptor NLR (TIR-NLR), G10-subclade CC NLR (CC_G10_-NLR), RESISTANCE TO POWDERY MILDEW 8-like CC NLR (CC_R_-NLR), and TIR-NB-ARC-like-β-propeller WD40/tetratricopeptide-like repeats (TNPs) (Kourelis et al., 2021). NLRs often function in networks, in which sensor NLRs recognise pathogen effectors and other NLRs function as helpers that translate the effector recognition into HR (Adachi et al., 2019). For example, CC_R_-NLRs are often considered TIR-NLR helpers, while MADA-containing CC-NLRs are often considered helpers of other CC-NLRs (Adachi et al., 2019; Derevnina et al., 2021; Contreras et al., 2023).

*Erysiphe pisi* employs a sophisticated array of effector proteins to disrupt host cellular processes and establish fungal colonisation. Recent studies have identified and characterised from 7 to 167 (Gupta et al., 2020, Sharma et al., 2019, respectively) putative *E. pisi* effectors with many different functional annotations, which are integral to pathogenicity and host defence evasion (Bhosle et al., 2019; Sharma et al., 2019; Bhosle & Makandar, 2021a). *Erysiphe pisi* effector expression varies according to the infection stage and the specific host (Gupta et al., 2020). Among the identified *E. pisi* effector candidates there are Egh16H homologs, ribotoxins/ribonucleases, glycoside hydrolases, and heat shock proteins (Sharma et al., 2019; Gupta et al., 2020). Bhosle et al. (2019) uncovered that three different *E. pisi* effectors were suppressed by the *er2* resistance gene in *P. sativum* accession JI-2480, highlighting the dynamic interplay between *E. pisi* effectors and host resistance mechanisms.

This work aimed to explore the transcriptomic networks involved in the interaction between *L. sativus* and *E. pisi* in resistant, partially resistant, partially susceptible accessions compared to susceptible accession. We used dual RNA-seq to reveal the key host resistance related genes (including NLRs), and pathogen effectors influencing this interaction, to deepen our understanding on the resistance mechanisms in *L. sativus* and the virulence strategies of *E. pisi*. This knowledge could significantly contribute to breeding programs in *L. sativus* and other legumes susceptible to *E. pisi*, such as pea.

## 2. Material and Methods

### 2.1. Plant material, growing conditions, and pathogen inoculation

Four contrasting *L. sativus* accessions were selected from a *L. sativus* worldwide collection previously phenotyped for the response against *E. pisi* using detached leaflets under controlled conditions (Martins et al., 2023): PI268478, PI221467_A, PI426882, and PI426890, rated as resistant (R), partially resistant (PR), partially susceptible (PS), and susceptible (S), respectively. Seedlings were grown in 0.5 L pots containing 250 cm^3^ of peat in a growth chamber at 20°C, 12-h light/12-h dark photoperiod. Fourteen-days old seedlings were inoculated with *E. pisi* isolate Ep-CO-01, which was permanently maintained on seedlings of the susceptible pea cv. ‘Messire’, at the Institute for Sustainable Agriculture – CSIC (Cordoba, Spain). The four *L. sativus* accessions were inoculated with *E. pisi* spores in five independent inoculation events corresponding to the five different inoculated time points (6, 12, 24, 48, 72 hours after inoculation (hai), using a settling tower to ensure a uniform conidial deposition of 8 conidia/mm^2^ onto each seedling. Leaflets from each plant were sampled at 0 (non-inoculated), 6, 12, 24, 48, and 72 hai and were immediately frozen in liquid nitrogen and stored at -80°C until RNA isolation. Time points 0, 12, 48, and 72 hai were selected for RNA-Seq analysis based on the different infection stages of *E. pisi* in the closest species *P. sativum* (Barilli et al., 2014). The experimental design is depicted in Fig. S1. Seven and fourteen days after inoculation, disease severity (DS) and infection type (IT) were visually estimated on the leaflets of *L. sativus* accessions and the pea cv. ‘Messire’ susceptibility control (Fig. S2). DS was scored as the percentage of leaflet area covered by mycelia. IT was recorded according to a 0 to 4 scale, where 0 corresponds to no visible disease symptoms and 4 corresponds to well-developed, freely sporulating colonies (Trabanco et al., 2012).

### 2.2. RNA isolation, library construction, and Illumina sequencing

For total RNA isolation, frozen leaflets were ground to a fine powder in liquid nitrogen using a mortar and pestle, and RNA was isolated using the GeneJET^TM^ Plant RNA Purification Mini Kit (ThermoScientific^TM^, Massachusetts, USA) according to the manufacturer’s instructions. RNA integrity and DNA contamination were assessed by electrophoresis in a 1.2% agarose gel stained with SYBR^TM^ Safe (Life Technologies^TM^, California, USA). Trace amounts of DNA contamination were removed from RNA with a treatment with TURBO^TM^ DNase (Invitrogen^TM^ by ThermoFisher Scientific^TM^, California, USA), following the manufacturer’s instructions. RNA concentration was measured using a Qubit 2.0 fluorometer with the Qubit RNA BR (Broad-Range) Assay Kit (Life Technologies^TM^, California, USA). RNA purity was estimated based on the 260/280 and 260/230 absorbance ratios using a NanoDrop^TM^ 2000c Spectrophotometer (Thermo Scientific^TM^, Passau, Germany). The RNA library construction was carried out using a Stranded mRNA Library Preparation Kit (Roche/KAPA mRNA HyperPrep kit) and samples were sequenced by lllumina Novaseq PE150 at STABvida sequencing provider (Lisbon, Portugal).

### 2.3. Bioinformatics analysis

For data analysis, all the reads from the sequencing data were subjected to a quality check using FastQC v0.11.9 (Andrews, 2010). Adaptor, barcodes, and low-quality reads (Phred score < 20) were removed using Cutadapt v4.0 (Martin, 2011) and high-quality reads were aligned to the *L. sativus* genome (JIC_Lsat_v2.1.1) (Vigouroux et al., 2024) using HISAT2 v2.2.1 with paired-end parameters (Kim et al., 2015) (Table S1). Read counts per gene were obtained using featureCounts v2.0.6 (Su et al., 2014) based on the corresponding genome annotation. To analyse the relationship among biological replicates and the differences among time points and accessions, a principal component analysis (PCA) was done using the normalised gene counts per sample. Differential expression analysis was performed using the DESeq2 R package by comparing the expression profile of the PS, PR or R accessions with that of the S accession, for each time point. False discovery rate (adjusted *P*-value) < 0.05 and |log_2_ fold change| > 1.0 were set as the thresholds for significant differential expression (Table S2).

Clustering analysis of differentially expressed genes (DEGs) based on expression patterns was performed using k-means clustering. Due to the limited functional annotation of the JIC_Lsat_v2.1.1 proteome, Mercator4 v6.0 (Lohse et al., 2014) was used to predict the functional annotation of each DEG identified. To increase the protein annotation rate, two additional tools were selected within Mercator: the annotation tool ProtScriber v0.1.3 and the BLAST tool, which provides Swiss-Prot protein annotations for similar proteins using the Swiss-Prot dataset of Viridiplantae proteins. To complement gene functional annotation of DEGs, candidate *A. thaliana* orthologues were identified using Blastp (v2.15) analysis based on *L. sativus* predicted protein sequences (Vigouroux et al., 2024). Only the best hits with an *e-*value < 0.001 were selected. Gene Ontology (GO) functional enrichment analysis of the DEGs was performed with g:Profiler (Kolberg et al., 2023) with Benjamini–Hochberg multiple testing correction (*P-*< 0.05) (Benjamini and Hochberg, 1995).

NLRtracker v1.3.1 (Kourelis et al., 2021) was used to predict complete nucleotide-binding leucine-rich repeats (NLRs) on the JIC_Lsat_v2.1.1 proteome. To accurately assign each NLR to a class, we aligned the RefPlantNLR dataset (Kourelis et al., 2021) together with the NB-ARC output file of NLRtracker using Clustal Omega v1.2.2 within Geneious Prime 2022.2.2 (Sievers & Higgins, 2014). A *L. sativus* NLR phylogenetic tree (Fig. S3) was obtained using the FastTree algorithm (Price et al., 2010) with the pseudo-counts setting, and the tree was rooted using non-plant NLRs from the RefPlantNLR dataset (Kourelis et al., 2021). The clades for each NLR class were identified based on phylogenetic clustering with the respective reference NLRs.

Heat maps were generated using the heatmaply R package. The expression values were an average of three biological replicates, except for the sample PR at 48 hai, which only pertained to a single biological replicate.

For *E. pisi* candidate effector prediction, the high-quality RNA-Seq reads from inoculated samples (12, 48 and 72 hai) were aligned to the *E. pisi* ASM20880v1 NCBI genome using HISAT2 2.2.1 (Kim et al., 2015). Uniquely mapped aligned reads (*E. pisi* specific) were extracted using samtools v1.9 (Li et al., 2009) and the merged left and right reads were used to build a *de novo* transcriptome using Trinity (Grabherr et al., 2011). This transcriptome was then used as reference to estimate gene expression by aligning again the RNA-seq reads using Salmon v1.10.2 (Patro et al., 2017).

*Erysiphe pisi* effector candidates were predicted in the *de novo* transcriptome’s open reading frames using Predector (Jones et al., 2021) where a minimum Predector score of 0 was used as the threshold. Effectors’ structures were predicted using ColabFold (Mirdita et al., 2022), and those with a predicted template modelling score higher than 0.5 were kept for further analysis. To search for effectors with structural similarities to known effector structures, we used Foldseek (van Kempen et al., 2023) on the well-predicted effector structures. We compared our predicted structures to a Foldseek database (https://zenodo.org/records/6480453), with 26,675 known effector structures of 21 species (Seong & Krasileva, 2023). On the other hand, the BLASTp tool was used against the NCBI fungi database for sequence-based effector functional prediction, using an e-value cut-off of 1E-05. The BLASTp tool was also used against fungi databases for functional prediction of the Foldseek hits (van Kempen et al., 2023).

### 2.4. Validation of RNA-Seq data using quantitative real-time PCR (RT-qPCR)

To validate RNA-Seq data and follow the whole infection response process trailed (0, 6, 12, 24, 48, and 72 hai) nine DEGs were selected: *inactive beta-amylase 9* (g5179), *kunitz trypsin inhibitor 5* (g3907), *NAC domain-containing protein JA2L* (g1304), *peptidyl-prolyl cis-trans isomerase FKBP65* (g28159), *eugenol synthase 1* (g18115), *pathogenesis-related protein 10* (g4535), *adagio protein 3* (g27034), *protein EARLY FLOWERING 4* (g19671), *glycine-rich RNA-binding protein 7* (g29906). These genes were selected based on their high expression variation on the different accessions and time points. cDNA was synthesised from 1 μg of total RNA from each sample following the manufacturer’s instructions of the iScript™ cDNA synthesis kit (Biorad, California, USA). For RNA-Seq validation by RT-qPCR, the comparisons were performed using the 0 hai time point (non-inoculated) as the reference to the corresponding accession.

Specific primers were designed using the Primer3Plus online tool (https://primer3plus.com/) (Boston, USA), and checked for specificity using the Primer-BLAST tool (NCBI, USA). Primers were designed in the 3′ intra-exonic regions and were synthesised by STAB Vida (Caparica, Portugal) (Table S3).

Four reference genes previously identified in *Lathyrus* species were selected to evaluate expression stability in the samples and conditions under study: β*-tubulin* (contig nr a6507;507), γ*-tubulin* (contig nr a77720;50), *histone H2A.2* (contig nr a20510;122), and *chromodomain helicase DNA-binding protein* (contig nr a1310;251) (Almeida, et al., 2015; Santos et al., 2018). The expression stability was tested using the geNorm (Vandesompele et al., 2002), NormFind (Andersen et al., 2004), BestKeeper (Pfaffl et al., 2004), the ΔCt method (Silver et al., 2006), and RefFinder (Xie et al., 2012) online tool (https://blooge.cn/RefFinder/).

The relative expression of the nine selected target genes was determined by quantitative real-time PCR (RT-qPCR). The RT-qPCR reactions were performed using three biological replicates per accession (S, PS, PR, R) and six time points (0, 6, 12, 24, 48, and 72 hai). RT-qPCR was performed in a final volume of 20 µl, containing 0.5 ng of cDNA, 0.5 µM of each primer (except for pathogenesis-related protein 10 where 1 µM was used), and 1 x LightCycler® 480 SYBR Green I Master. Thermal cycling for target and reference genes started with a denaturation step at 95 °C for 5 min, followed by 40 cycles of denaturation at 95°C for 10 s and 60°C for 30 s. At the end of all gene expression cycling protocols, melting curve analysis was performed to validate amplification specificity under the following conditions: 65°C for 1 min to 97°C with the increment of 0.5°C for 11 s. Also, a negative template control (NTC) without cDNA was included in each PCR plate to detect possible genomic DNA contaminations. The relative expression values (fold change-FC) of the nine target genes were normalised to the non-inoculated samples (0 hai) and the two reference genes showing the highest expression stability using the Pfaffl method (-Efficiency ^ΔΔCt^) (Pfaffl, 2001). Finally, FC data were transformed into a logarithmic scale (base 2) for graphical representation and statistical analyses. ANOVA, followed by Dunnett’s multiple comparisons test, was performed to compare the expression levels of each time point to the non-inoculated sample per accession. Linear regression was performed to assess the relationship, and Pearson’s correlation test was used to evaluate the correlation between the log2 FC values of RNA-seq and RT-qPCR. The data was analysed using R statistical software version 4.3.0 (R Core Team, 2022) and GraphPad Prism 6 (GraphPad Software Inc.; San Diego, CA, USA).

## 3. Results

### 3.1. Dual RNA-Seq Analysis and RNA-seq validation by RT-qPCR

The selected *L. sativus* accessions (S, PS, PR, R) presented different whole plant disease severities (DS) at 7 and 14 days after inoculation (dai), as reported for detached leaflet assays (Martins et al., 2023) (Fig. S2). Using the Trabanco et al. (2012) disease symptoms visual scale, we observed a moderate mycelial development without sporulation or necrosis for all accessions at 7 dai, and for R and PR accessions at 14 dai (IT=3) (Fig. S2). At 14 dai, we observed an abundant mycelial development and profuse sporulation for S and PS (IT=4). By 14 dai, DS had increased for all accessions, although macroscopic sporulation was more evident in R and PR than in PS and S, which reached a comparable DS to pea cv. ‘Messire’ at 7 dai (Fig. S2). A total of 475,821,117 clean reads were generated for the four accessions across four time points, averaging 10.343.937 reads per sample. On average, 86.46% of sample reads were mapped to the JIC Lsat v2.1.1 *L. sativus* reference genome (77.33% up to 90.05%), while 0.48% of the infected sample reads mapped to the ASM20880v1 *E. pisi* genome (Table S1).

The gene expression PCA of the contrasting *L. sativus* accessions to *E. pisi* showed that the three biological replicates at each time point clustered closely (Fig. 1A). Furthermore, *L. sativus* samples revealed distinct gene expression patterns based primarily on accession (Fig. 1A). Looking at the first two principal components, both S and PS samples clustered apart from the rest of the samples, separating from R and PR on PC1 and distinguishing between themselves on PC2. On the other hand, R and PR accessions clustered together with slight differences mostly on PC2, showing a more similar transcriptional response to *E. pisi* infection. When focusing on the separation per time point within each accession, there were roughly two groups. Samples from 0 hai and 48 hai clustered closer together in all accessions, while samples at 12 and 72 hai formed a separate group.

**Fig. 1.**
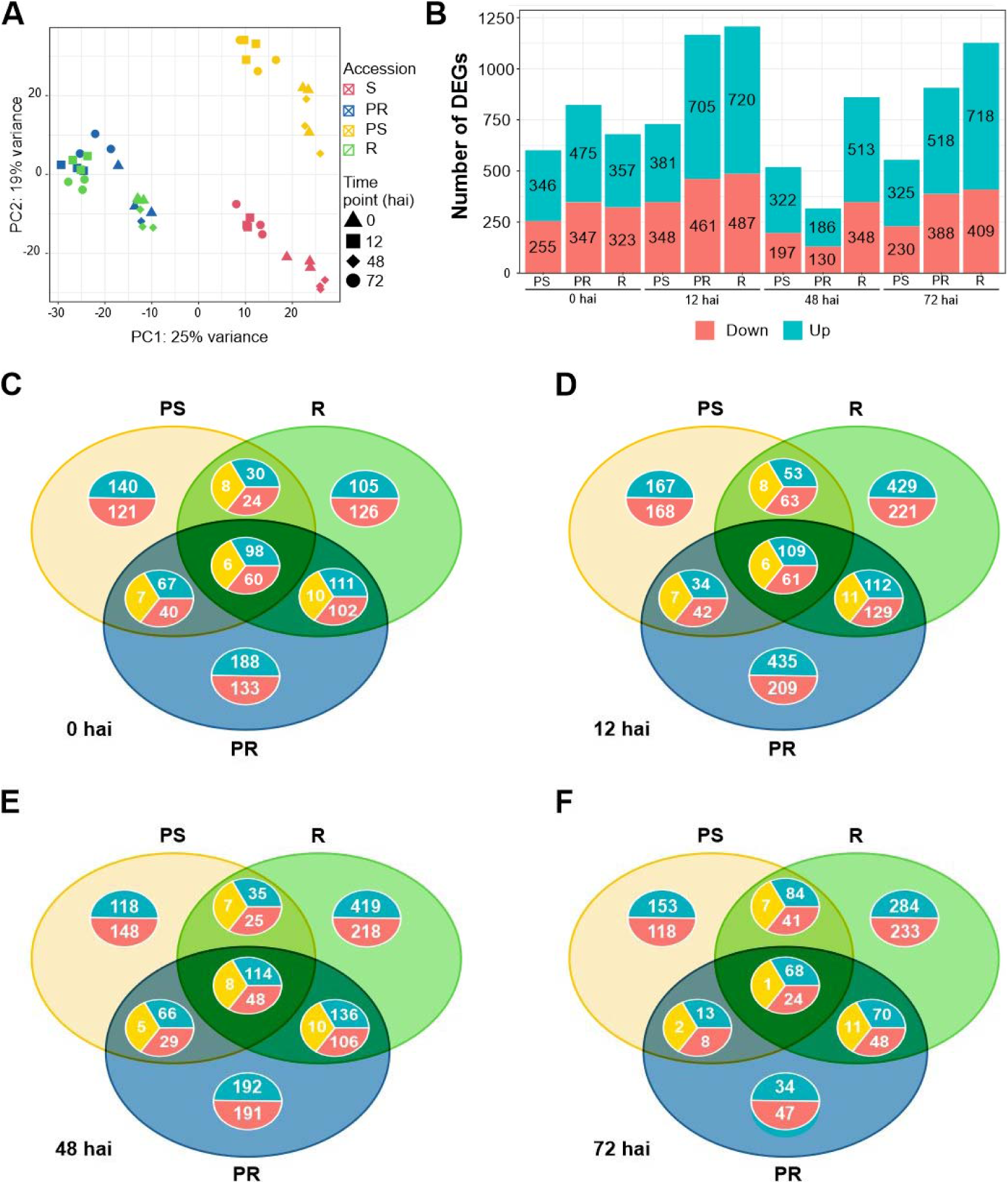
Overview of transcriptome data and differentially expressed genes (DEGs) in the *Lathyrus sativus* response to *Erysiphe pisi*. A) Principal component analysis of the time-series transcriptomes of the four contrasting accessions to *E. pisi* based on counts per million values of all genes. B) Number of DEGs upregulated (green) and downregulated (yellow) for comparisons under study. C-F) Venn diagrams showing unique and common DEGs between *L. sativus* accessions for each time point: 0 hai (C), 12 hai (D), 48 hai (E) and 72 hai (F). R – resistant, PR – partially resistant, PS – partially susceptible.

We identified a total of 3,109 DEGs, finding a larger number of upregulated genes than downregulated genes in all comparisons under study (Fig. 1B). DEGs commonly identified at 0 hai (non-inoculated conditions) and at least one inoculated comparison were selected to represent the basal defence response of *L. sativus* to *E. pisi*. The largest number of DEGs was identified for the R compared to S at 12 hai (720 upregulated and 487 downregulated), followed by the PR compared to S at 12 hai (429 upregulated and 293 downregulated) (Fig. 1B). At 12 hai, *E. pisi* triggered large transcriptional changes across all accessions, but by 48 hai, the number of DEGs had substantially decreased in all comparisons. The number of common DEGs across all accessions remained relatively stable over time (164, 181, and 170 for 0, 12, and 48 hai, respectively). However, at 72 hai, only 93 common DEGs were detected (Fig. 1C-F). Notably, transcriptional changes at 12 and 72 hai progressively increased from PS to PR and R accessions (Fig. 1D and F).

We observed a high Pearson’s correlation (r=0.98) between the log fold change (logFC) values obtained by RNA-seq and RT-qPCR for the nine genes tested. Additionally, linear regressions were fitted showing a coefficient of determination (R^2^) of 0.97 (Fig. S4A). Looking at the comparative heatmap with both RT-qPCR and RNA-seq data most gene expression patterns seemed analogous between RT-qPCR and RNA-seq throughout the *E. pisi* infection in all accessions (Fig. S4B), technically validating our RNA-seq datasets.

### 3.2. Constitutive transcriptional differences among *Lathyrus sativus* accessions

At non-inoculated conditions (0 hai), the PR is the accession with the most distinct *E. pisi* defence-related transcriptome compared to the S accession, with 475 upregulated and 347 downregulated DEGs (Fig. 1B, C). Notably, 321 DEGs were exclusive to the PR compared to S (188 upregulated, 133 downregulated) (Fig. 1C, Table S2).

Enrichment analysis revealed that PR-exclusive upregulated biological processes (BPs) included regulation of defence response (GO:0031347), response to jasmonic acid (GO:0009753), and defence response to other organisms (GO:0051707), among other defence-related processes (Table S4). In these BPs, defence-related DEGs with high logFC included *disease resistance protein RUN1* (g29548.t1, logFC=5.6) and two *Bowman-Birk type proteinase inhibitors* (g11928.t1 and g11927.t1, logFC=5.0 and 3.5).

In the R accession, unique and upregulated DEGs showed an enrichment in the phenylpropanoid and lignin biosynthetic processes (GO:0009809, GO:0009699). In contrast, the PS accession displayed a distinct constitutive response compared to R and PR accessions, with upregulated DEGs enriched in BPs related to general cellular and organismal responses to environmental stimuli. This includes response to stress (GO:0006950), and processes related to nitrogen metabolism, such as nitrate assimilation (GO:0042128) and the nitrogen cycle (GO:0071941) (Table S2 and 4).

At 0 hai, a total of 164 DEGs were commonly identified among the PS, PR, and R vs. S accession. These include 98 common upregulated DEGs, 60 common downregulated DEGs, and 6 DEGs showing ambiguous expression, being up or downregulated depending on the accession (Fig. 1C). Although functional enrichment did not reveal specific BPs for these common DEGs, several defence-related DEGs were observed, such as the NLR *putative disease resistance RPP13-like protein 1* (g14573.t1), which exhibited logFC > 4 across all accessions (Table S2 and 4).

### 3.3. Common and unique *Lathyrus sativus* transcriptional responses against *Erysiphe pisi*

Common DEGs among R, PR, and PS compared to S were predominantly upregulated in all inoculated time points, with enrichment in BPs related to stress and/or stimulus response (GO:0006950, GO:0050896). Upregulated defence-related DEGs included: *probable mannitol dehydrogenase,* (g8997.t1, g8994.t1, g8995.t1 logFC ranging from 5.4 to 8.2); BURP domain protein *RD22* (g22981.t1, g22990.t1, g22980.t1, g22989.t1, g22988.t1, g22982.t1, logFC ranging from 1.1 to 4.9); *peptidyl-prolyl cis-trans isomerase FKBP62* and *FKBP65* (g22345.t1, g13946.t1, g28159.t1, logFC ranging from 1.0 to 6.0); *Bowman-Birk type proteinase inhibitor* (g11928.t1, g11927.t1, logFC ranging from 1.7 to 5.0); and *polygalacturonase inhibitors* (g10621.t1, g11046.t1, g28453.t1, logFC ranging from 1.0 and 6.2). Common downregulated BPs across accessions were identified only at 12 hai and relate to starch (GO:0005982, GO:0019252) and glucan (GO:0009250) biosynthetic processes (Table S4). Overall, the number of downregulated defence-related DEGs was significantly lower compared to the upregulated ones.

Accession-specific DEGs revealed different response strategies among accessions. In the R accession, exclusive upregulated BPs at 12 hai were predominantly linked to physical and chemical defence barriers against pathogen infection. These included cell wall organisation and biogenesis (GO:0071555, GO:0071554), lignin biosynthetic process (GO:0009809), cutin biosynthetic process/cutin-based cuticle development (GO:0010143, GO:0160062), phenylpropanoid, polyamine and spermidine metabolic processes (GO:0009699, GO:0009698, GO:0008295, GO:0006595). Cell wall organisation and biogenesis processes (GO:0071555, GO:0071554) continued to be enriched at 48 hai in the R accession (Table S4).

At later infection stages (72 hai), the R accession showed upregulated BPs associated with protein folding (GO:0006457), response to osmotic stress (GO:0006970), oxidative stress and detoxification (GO:0006979, GO:0098754), and abscisic acid (ABA) response (GO:0009737). DEGs with putative antifungal functions were exclusively upregulated in R at 12 and/or 72 hai, including *eugenol synthase 1* (g18115.t1, logFC > 3.0) and *thaumatin-like protein 1* (g29711.t1, logFC = 2.9). Additionally, *hypersensitive-induced response protein 1* (g2782.t1, logFC > 1.4) was exclusively upregulated in R across all time points, including non-inoculated conditions (Table S2 and 4).

The BPs previously identified for exclusively upregulated DEGs in PR compared to S at non-inoculated conditions-0 hai (such as the BP related to response to biotic stimulus GO:0043207, GO:0051707, GO:0044419, GO:0009607,GO:0098542) were also observed at 12 hai, with some of them showing increased expression at this later time point (Table S4). However, at 48 and 72 hai, no specific GO term enrichment was identified for PR. Nevertheless, some defence-related DEGs, such as *Kunitz-type trypsin inhibitor-like 2* and *5* (g3757.t1, g3907.t1) were detected in these time points (Table S2).

At 12 hai, PR DEGs showed significant enrichment in BPs related to organismal responses to biotic stimuli and regulatory mechanisms of defence and stress responses. Exclusively downregulated DEGs in PR accession identified at 48 and 72 hai were associated with glycerol transmembrane transport (GO:0015793), polyol transmembrane transport (GO:0015791), carbohydrate catabolic process (GO:0016052), and starch metabolic and biosynthetic processes (GO:0005982, GO:0019252) (Table S4).

A larger number of DEGs were common between R and PR compared to S than between R and PS or PR and PS compared to S, except at 72 hai, where PR and PS compared to S shared more DEGs (132) than R and PR compared to S (119) (Fig. 1C-F, Table S2). In general, R and PR shared upregulated DEGs related to plant immunity, such as *receptor-like protein Cf-9 homolog* (g8326); cell wall organisation, *including 4-coumarate-CoA ligase CCL1* (g17573); antifungal activity, like *kunitz-type trypsin inhibitor-like 2* and *5* (g3757.t1, g3907.t1); and secondary metabolism genes, including *benzyl alcohol O-benzoyltransferases* (g9735, g9738). R and PR accessions also shared genes involved in jasmonic acid signalling, such as the *allene oxide cyclase* (g13281.t1, g13279.t1, g979.t1). At 12 hai (and at 0 hai), R and PR accessions shared upregulated DEGs enriched in trichome morphogenesis and differentiation (GO:0010026, GO:0010090), as well as plant epidermis morphogenesis (GO:0090626), including DEGs such as *Protein CPR-5* (g20886.t1) and *Protein SCAR2* (g12176.t1 and g12209.t1). At 72 hai, these DEGs were also shared between R and PS. At 48 hai, both resistant accessions (R and PR) had upregulated DEGs involved in responses to abiotic stresses, including reactions to inorganic substances, chemicals, oxygen-containing compounds, and general environmental stressors. Additionally, at both 48 and 72 hai, R and PR shared upregulated DEGs related to responses to water stress conditions, such as *NAC domain-containing protein JA2L* (g14943.t1 and g1304.t1). No BPs from downregulated genes/proteins were shared between R and PR compared to S (Table S2 and S4).

Regarding the exclusive DEGs in PS compared to S at 12 and 48 hai, the upregulated group was predominantly associated with responses to environmental stimuli, such as chemicals (GO:0042221), and oxygen level changes (GO:0001666, GO:0036293, GO:0070482). Additionally, we found upregulated DEG terms associated with defence mechanisms and responses to external biotic interactions both in PS compared to S only at 48 hai, and in PR compared to S at 12 hai. At 72 hai, no BPs were enriched for upregulated DEGs. (Table S2).

### 3.4. Temporal gene expression dynamics in *Lathyrus sativus* response to *Erysiphe pisi*

From the three independent K-means clustering analysis (Fig. 2, Table S5), we detected specific DEG clusters exhibiting a clear differential pattern along infection (Fig. 2). Regarding the R accession, it was notable that clusters 7, 9, and 10 contained consistently upregulated DEGs in inoculated conditions compared to 0 hai (Fig. 2C). Common BPs found in these three clusters included response to chemicals, stimulus, and stress. In cluster 7, DEGs were mainly involved in response to oxidative stress (GO:0006979, GO:0098754, GO:0042744), such as five genes annotated as *peroxidase 4* (g20878.t2, g20880.t1, g1197.t1, g8235.t1, g1192.t1). *Peroxidase 4* DEGs were also upregulated in R compared to S, especially at 72 hai (Table S2). Two exclusive DEGs within cluster 9 (*S-adenosylmethionine decarboxylase proenzyme* (g13308.t1) and *tryptophan synthase beta chain 1* (g3000.t1)) were linked to GO terms describing processes related to the biosynthesis and metabolism of polyamines (GO:0006596, GO:0006595), biogenic amines (GO:0042401, GO:0009309, GO:0006576) and spermidine (GO:0008295, GO:0008216). Cluster 10 contained DEGs that showed the most pronounced upregulation after inoculation, particularly at 72 hai (Fig. 2C). In this cluster, DEGs were enriched for GO terms related to cellular responses to changes in oxygen levels (GO:0001666, GO:0036293, GO:0070482, GO:0071456, GO:0036294, GO:0071453, GO:1901700), to heat and temperature stimuli (GO:0009408, GO:0009266), as well as general stress responses such as protein folding (GO:0006457). Notable examples include class II *heat shock proteins* that respond to both temperature and oxygen level changes (g21471.t1, g21472.t1, g14212.t1, g22760.t1, g22740.t1), as well as other *heat shock proteins* that are generally upregulated in R compared to S (Table S2 and S6). Notably, in cluster 10, DEGs were more highly expressed in the S accession at 0, 12, and 48 hai compared to R, indicating that BPs related to cellular responses to changes in oxygen levels, heat and temperature stimuli, and protein folding (Table S6) were more important for resistance to *E. pisi* at 72 hai (Table S2).

**Fig. 2.**
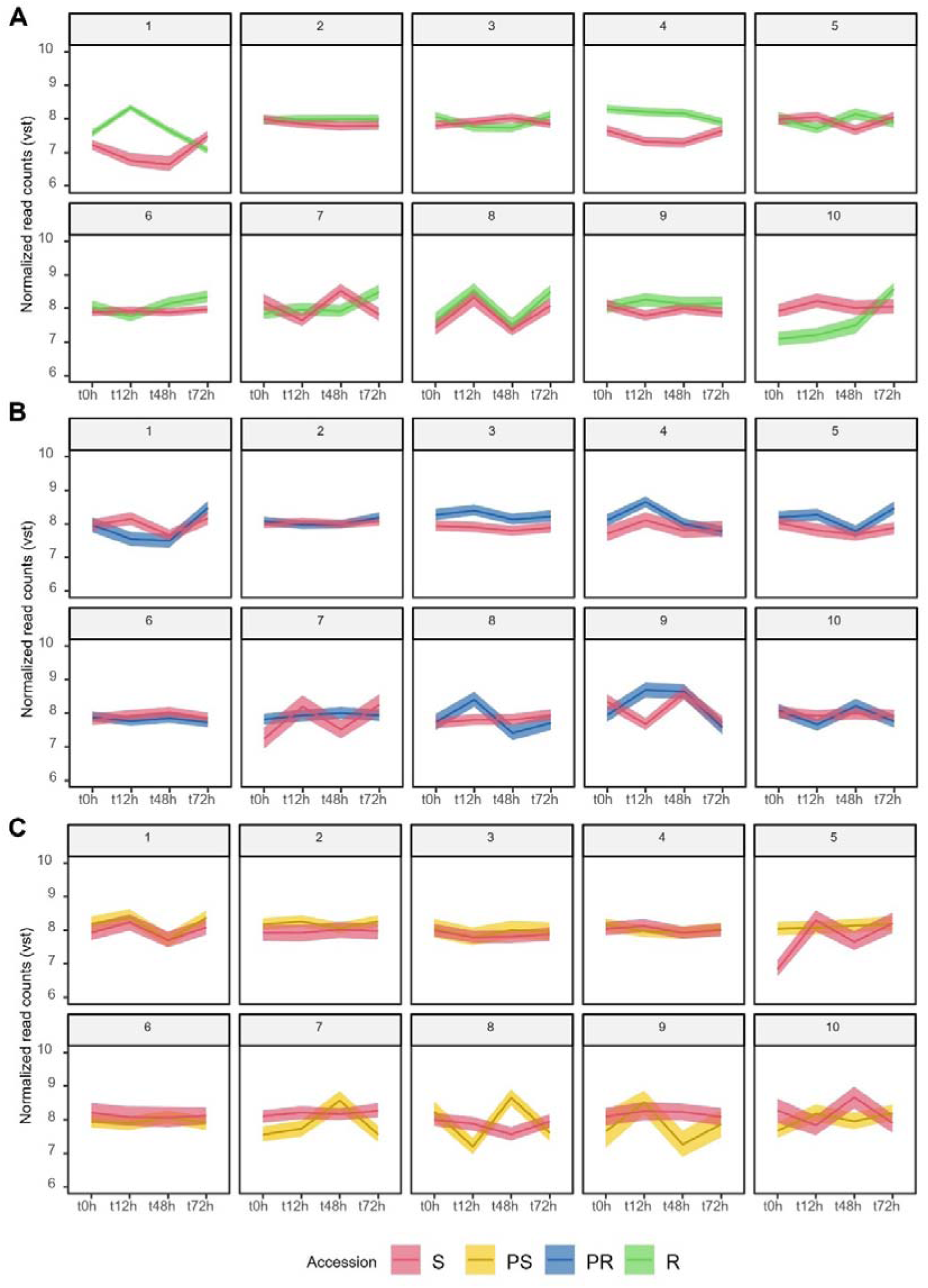
– Grouping of temporal gene expression dynamics of *Lathyrus sativus* after *Erysiphe pisi* infection by K-means clustering of vst-normalised DEGs. (A) R and S, (B) PR and S, (C) PS and S. R – resistant, PR – partially resistant, PS – partially susceptible, S – susceptible.

In contrast to clusters 7, 9, and 10, where the R accession DEGs were upregulated, cluster 4 contained DEGs that were downregulated at 12, 48, and 72 hai compared to 0 hai, although the degree of downregulation was relatively low (Fig. 2C). BPs identified included the biosynthesis, metabolism, and catabolism of lignin and phenylpropanoids, alongside their roles in cell wall organisation, carbohydrate metabolism, and stomatal movement regulation (Table S6).

The expression dynamics of R DEGs in clusters 1 and 6 showed distinct patterns: in cluster 1, DEGs were highly expressed during the early response to *E. pisi* infection (12 hai), with expression levels decreasing at later time points (48 and 72 hai). In contrast, cluster 6 exhibited the opposite trend, where DEGs had a lower expression at 12 hai but an increase in expression at 48 and 72 hai (Fig. 2C). DEGs more expressed at 12 hai (cluster 1) are mainly involved in DNA replication, confirmation and unwinding (GO:0006268, GO:0071103, GO:0006270, GO:0032392, GO:0032508), chromosome organisation (GO:0051276), cutin and cuticle biosynthesis/development (GO:0160062, GO:0010143), and cell wall organisation or biogenesis (GO:0071554, GO:0071555). Conversely, the BPs that were more prominently regulated during the later stages of infection (cluster 6) included the response to hormones (GO:0009725) and hormone-mediated signalling pathways (GO:0009755), in addition to the general response to stress and stimuli. Examples of DEGs enriched in these BPs are: *WRKY transcription factors* (g20902.t1, g12139.t1), *ethylene-responsive transcription factors* (g1394.t1, g6994.t1, g8619.t1, g19521.t1, g22398.t1), *NAC domain-containing proteins 21/22* (g1645.t1), *protein TIFY 10B* (g4147.t1*), peptidyl-prolyl cis-trans isomerase CYP40* (g28974.t1). This cluster also encompassed responses to jasmonic acid, including signalling pathways (GO:0009753, GO:0009867, GO:0071395), responses to endogenous stimuli (GO:0009719), and responses to varying oxygen levels (GO:1901700, GO:0071456, GO:0071453, GO:0036294) (Table S2 and S6).

In the PR accession, clusters 3, 4, and 8 contained DEGs that may play crucial roles in early response stages (12 hai) but decrease their expression at 48 and 72 hai, even when compared to 0 hai (Fig. 2B). Many of the DEGs in clusters 3 and 8 were functionally enriched for response to other organisms (GO:0051707, GO:0044419), and response to biotic stimulus (GO:0009607, GO:0043207). This is also the case of cluster 9, where DEGs involved in these BPs were also upregulated at 48 hai (Fig. 2B). Most of the BPs enriched in cluster 4 were shared with clusters 3, 8, and 9, covering responses to stress (GO:0006950), response to stimulus (GO:0050896), response to chemicals (GO:0042221), and response to abiotic stimulus (GO:0009628). Cluster 6 included DEGs that exhibited reduced expression at all inoculated time points compared to the 0 hai condition and to the expression profile in S (Fig. 2B). These DEGs were enriched in general GO terms related to the response to stimulus (GO:0050896, GO:0006950), and more specifically in negative regulation of SAR (GO:0010113), which include *E3 SUMO-protein ligase SIZ1* (g270.t1), *UDP-glycosyltransferase 71K2* (g8457.t1 and g8456.t1), and *inorganic pyrophosphatase TTM2* (g15582.t1).

In the PS accession, clusters 4 and 5 were primarily associated with general stress and stimulus responses, with expression levels similar between non-inoculated and inoculated conditions (Fig. 2A). An example of a gene involved in these responses is *linoleate 9S-lipoxygenase* (g14089.t1) in cluster 4. Conversely, DEGs enriched for stimulus and hormone responses in cluster 8 were less expressed at 12 and 72 hai than at 0 and 48 hai (Fig. 2A). These include DEGs such as *ethylene-responsive transcription factors* (ERF23/34, g31009.t1, g13966.t1). Lastly, in cluster 10, BPs related to secondary metabolism, including the phenylpropanoid biosynthetic process (GO:0009699) and the (-)-pinoresinol biosynthetic and metabolic processes (GO:1901599 and GO:1901598), were upregulated following inoculation, particularly at 12 and 72 hai. An example of a gene involved in these processes is the *disease resistance response protein 206* (g27378.t1, g27379.t1, g27377.t1), which plays a role in the (-)-pinoresinol biosynthetic and metabolic pathways (Table S2 and S6). Overall, the S accession exhibited more similar and overlapping patterns with PS (clusters 1, 2, 3, 4, and 6) than with PR (clusters 2 and 6) or R (clusters 2 and 8) accessions, patterns that were overall stable from 0 to 72 hai, except for cluster 1 in PS and cluster 7 in R Fig. 2 A-C, red lines.

To further investigate the gene expression dynamics during *E. pisi* infection, we quantified by RT-qPCR the expression of selected defence-related DEGs in two additional time points: 6 hai and 24 hai (Fig.S5). The expression of *FKBP65* (upregulated in R, PR, and PS compared to S, (Table S2) was consistent across all accessions, with higher expression at 12 hai, followed by a decrease in expression reaching lower expression levels at 24 or 48 hai and then a secondary increase at 72 hai especially in R where expression spiked at 72 hai (Fig.S5).

The Kunitz trypsin inhibitor 5 was upregulated in R and PR compared to S (Table S2) and displayed similar expression trends between S and R accessions, with S showing lower expression levels. PS and PR accessions also shared similar expression profiles. Notably, at 72 hai, expression levels decreased in S and PR but increased in PS and R (Fig.S5)

*NAC domain-containing protein JA2L* (upregulated at 48 and 72 hai in R and PR compared to S) was barely expressed in S and PS accessions but showed a consistently increasing expression in R across all time points (Fig.S5). *Eugenol synthase 1* was not expressed in the PS accession but followed the same expression pattern in R, S, and PR, with higher expression at 12 and 72 hai. PR showed a notably higher expression level compared to R and S at 72 hai. R was the only accession with early *synthase 1* early expression at 6 hai (Fig.S5).

*Pathogenesis-related protein 10* showed a gradual increase in expression over time, with the highest levels observed in R (Fig.S5).Finally, *glycine-rich RNA-binding protein 7* exhibited a similar expression profile in S, PS, and PR accessions, with a decrease until 12 hai, an increase up to 48 hai, followed by another decrease at 72 hai. In R, expression remained steady during the first 24 hours, with a slight decline at 48 and 72 hai (Fig.S5).

### 3.5. Expression of *Lathyrus sativus* NLR genes during *Erysiphe pisi* infection

Among the 3,109 DEGs detected when comparing the R, PR and PS accessions to the S within each time point, we identified a total of 52 NLR genes: 22 CC-NLRs, 25 TIR-NLRs, 3 CC_G10_-NLRs, and 2 CC_R_-NLRs (Table S2).Analysing the normalized read counts of each identified NLR, we observed the formation of roughly three distinct groups (Fig. 3). The blue (first) and the pink (third) groups mostly depicted highly expressed NLRs found in all samples, although with punctual fluctuations in expression per time point. The second cluster mainly contained accession-specific NLRs, with low expression in some accessions.

**Fig. 3.**
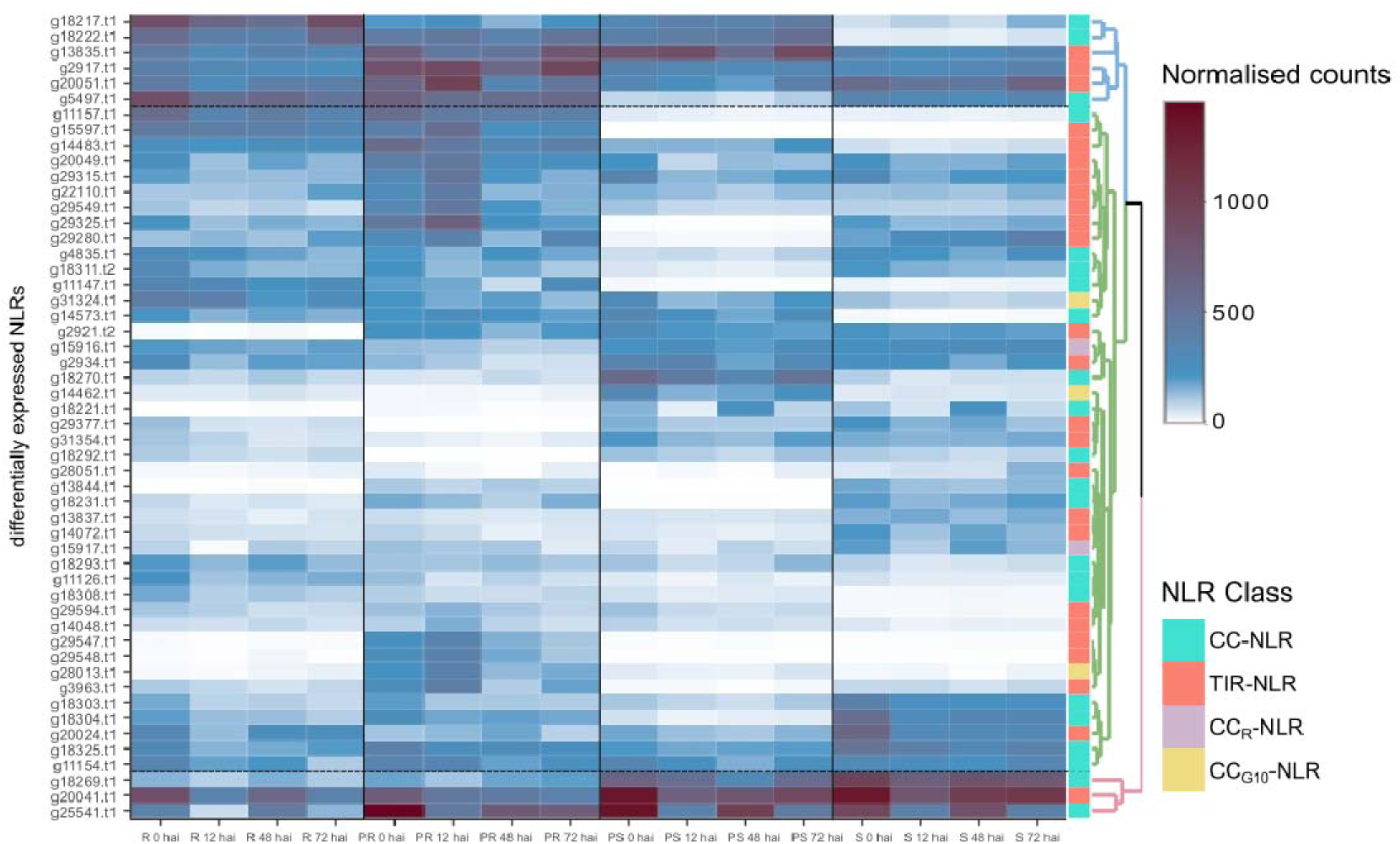
Expression heatmap of differentially expressed NLRs in four contrasting *Lathyrus sativus* accessions at 0, 12, 48 and 72 hours after *Erysiphe pisi* inoculation. NLR classes are represented in the last column with the following colour code: CC-NLR – turquoise; TIR-NLR – salmon; CC_R_-NLR – lilac; CC_G10_-NLR - yellow. K-means clustering clustered NLR expression patterns in three groups visible on the dendrogram on the right side (pink, green and blue). R – resistant, PR – partially resistant, PS – partially susceptible, S – susceptible.

Sixteen NLRs were highly expressed in all accessions, with at least 100 average normalised counts (showed by light blue) in one of the time points (Fig. 4). These included: six *RUN1* homologues (g2917.t1, g13835.t1, g20024.t1, g20051.t1, g22110.t1, and g29549.t1), three *RPP13-like proteins* 1 (g18269.t1, g18303.t1, g18325.t1), *RGA2* (g11154.t1), *SUMM2* (g31324.t1) and a homologue of the *At5g66900* gene also known as N-requirement gene 1 (*NRG1.1*) (g15916.t1).

**Fig. 4.**
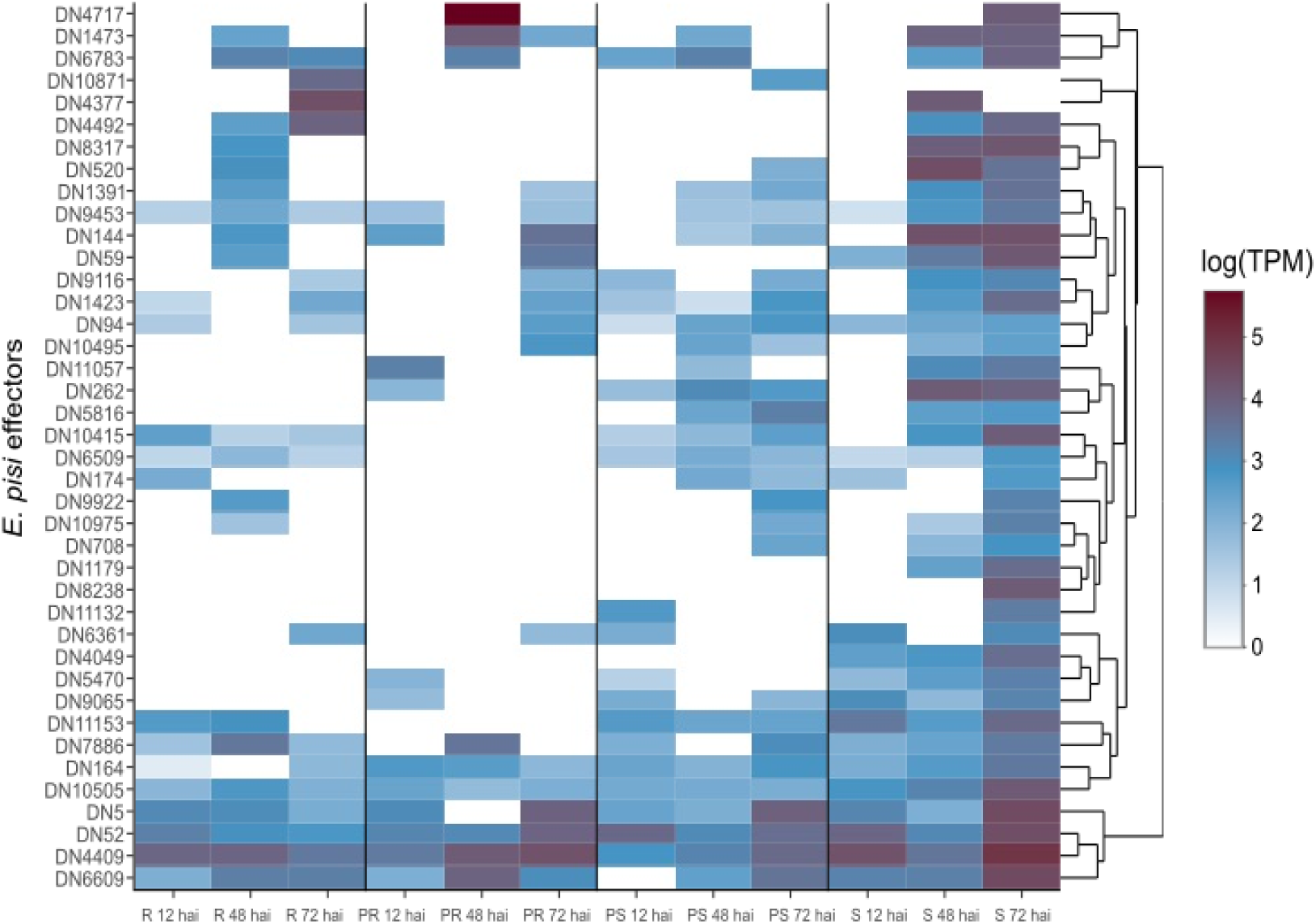
Expression heatmap (in transcripts per million) of potential *Erysiphe pisi* effectors at 12, 48 and 72 hours after inoculation (hai) in *Lathyrus sativus* accessions with contrasting powdery mildew responses. R – resistant; PR – partially resistant; PS – partially susceptible; S – susceptible.

The time point with the highest number of upregulated differentially expressed NLRs was 12 hai (23), followed by 0 hai (21), 72 hai (16), and 48 hai (12) (Fig. 4, Table S2). In contrast, 72 hai was the time point with more downregulated NLRs (21), followed by 0 and 12 hai (19), and finally by 48 hai (15) (Fig. 4, Table S2). PR was generally the most different accession from S when it came to NLR expression, with 21 differentially upregulated NLRs across time points, while R had 13 upregulated NLRs and PS only eight (Table S2, Fig. 4). At 12 hai, PR accession showed the largest amount of upregulated NLRs (18), followed by PR at 0 hai (15), R at 0 hai (12), PR at 72 hai (11), and R at both 12 and 48 hai (10) (Table S2, Fig. 3).

Most differentially expressed NLRs (39/52) belonged to the CC- and TIR-NLRs: *RUN1* (18), *RPP13-like proteins 1* (12), *RGA* genes (5), *RPM1* (4). *RUN1* TIR-NLRs were found upregulated in all accessions and time points, especially in PR and at 12 hai samples, and some were also notably downregulated (g29280.t1, g29377.t1) (Fig. 3, Table S2). *RUN1* g15597.t1 was highly upregulated for R and PR accessions (Fig. 3). *RPP13-like proteins 1* CC-NLRs were also differentially expressed throughout accessions and time points, with g14573.t1 having notably high logFC in all accessions (Table S2). *RGA1* CC-NLR (g11157.t1) was highly expressed in R and PR accessions only, across all time points (Fig. 3). *RGA2* (g11154.t1) was only downregulated for R at 72 hai (Table S2). *RGA3* homologue g11147.t1 behaved similarly to *RGA1*, but g11126.t1 upregulation was R-specific at 0 and 72 hai (Table S2). *RPM1* CC-NLR g18222.t1 was highly upregulated in all samples, while g18217.t1 was upregulated in all R and PS samples (Table S2). *RPM1* homologues g18221.t1 and g18231.t1 were downregulated: g18221.t1 in PR and R, g18231.t1 for all PS samples and R at 72 hai (Table S2).

Additionally, we also identified nine differentially expressed NLRs that did not belong to any of the previous homolog groups. Such was the case for *SUMM2* (g31324.t1), upregulated at 0, 12 and 48 hai in R samples; *At4g27190* (g14462.t1) upregulated at 0, 48 and 72 hai samples in the PS accession; two *NRG1.1* homologues downregulated, g15916.t1 at 72 hai in the PR accession, and g15917.t1 at 12 hai in the R accession.

### 3.6. *Erysiphe pisi* effector candidates exhibit infection-stage specific expression patterns

A total of 2,874,699 reads mapped against the *de novo E. pisi* transcriptome, from which 1,660,268 reads were uniquely mapped. We then performed effector prediction on this transcriptome to further analyse the expression dynamics of effectors and link their impact on host differential expression. We identified a total of 297 potential effector proteins, of which 163 (Table S7) had a predicted template modelling score above the threshold. From these, 40 showed structural similarity to known fungal effectors from the Foldseek database (Table S7). The most common species to have a structurally similar protein to the *E. pisi* effector candidates were *Puccinia graminis*, *Magnaporthe oryzae, Rhizoctonia solani*, *Phytophthora infestans*, and the cereal powdery mildew causal agent *Blumeria graminis* (Table S8).

Among 40 effectors identified, the majority were categorised as secreted proteins, cell surface proteins, enzymes with varied functions, and some had sequence similarity to well-studied effectors from other species, such as an homologue of the *Magnaporthe oryzae SnodProt1* (Table S8). From the 40 *E. pisi* effector, 35 candidates exhibited sequence similarity to other proteins from the *Erysiphe* genus (Table S8).

By analysing the read counts of the predicted effectors across the infected samples, we observed that the S and PS accessions showed a consistent increase in the number of expressed effectors over time, rising from 15 to 38 in S, and from 17 to 26 in PS (Fig. 4). In contrast, R and PR accessions kept the number of expressed effectors lower than in S and PS accessions. In the R accession, 13 effectors were expressed at 12 hai, increasing to 20 at 48 hai, but then decreasing to 18 at 72 hai. In the PR accession, 12 effectors were expressed at 12 hai, 9 effectors expressed at 48 hai, and 16 effectors expressed at 72 hai (Fig. 4).

From the 40 effector candidates (Fig. 4; Table S7), 16 were found expressed in all accessions, including two secreted effector proteins (DN1391, DN144), a 7-dehydrochoslesterol reductase (DN10505), an alpha-glucosidase (DN9453), the SnodProt1 (DN1473), and a subtilisin-like protease 2 (DN7886). Among the expressed effector candidates, we found a pectin lyase-like protein (DN174) and a secreted glycosidase (DN9922) both expressed in R, PS and S samples in different time points (Table S7). Three effectors were only present in PS and S: a bacterial alpha-l-rhamnosidase domain protein (DN11132), a serine-threonine protein phosphatase (DN708), and a V-type proton ATPase subunit C (DN5816). Three effectors were exclusively found in the S accession: a fungal-specific transcription factor domain-containing protein (DN8238), a histone acetyltransferase ELP3 (DN4049), and a pre-rRNA processing protein (DN1179). The number of expressed *E. pisi* effectors was maximum at 72 hai in the S accession, where 38 out of 40 effector candidates were found expressed (Fig. 4; Table S7).

## 4. Discussion

In this study, we employed a dual transcriptomics approach to gain deeper insights into the molecular interactions of four *L. sativus* accessions exhibiting contrasting responses to *E. pisi*, during the initial stages of powdery mildew infection. The transcriptional profiles highlighted how diverse *L. sativus* genetic backgrounds modulate their interaction with *E. pisi*, revealing distinct defence mechanisms specific to each accession, while also identifying key common defence responses. Additionally, we analysed the expression of defence-related NLR genes across all accessions and time points, finding that NLR expression depends on the plant genetic background. Among these NLRs, we found differences for gene homologues which could be involved in powdery mildew resistance, such as *RUN1*, *RPP13-like protein 1*, two *RPM1* proteins, and an *RGA3*. On the pathogen side, we detected putative *E. pisi* effectors that employ different virulence strategies. While some effectors were already described, most belonged to broad and previously undescribed categories in *E. pisi*, further expanding our knowledge on the infection mechanisms of this pathogen.

*Erysiphe pisi* triggered a biphasic *L. sativus* response, characterised by an initial burst in differential gene expression at 12 hai, followed by a quiescent phase at 48 hai, during which the pathogen continued its development but triggered much lower levels of differential gene expression. A second wave of intense gene expression was observed at 72 hai. This trend was consistently observed when additional time points were included for RT-qPCR analysis of selected genes, with expression at 6 hai and 24 hai mirroring that at 12 hai and 48 hai, respectively. A similar biphasic response has also been documented in soybean (*Glycine max*) infected by rust (*Phakopsora pachyrhizi)*, suggesting a conserved defence strategy across legume species when facing aerial diseases caused by fungal pathogens which rely on haustoria formation (van de Mortel et al., 2007; Schneider et al., 2011).

On the pathogen side, we noted that effector expression increased over time, peaking at 72 hai, a trend also observed for the powdery mildew agent *Blumeria graminis* f. sp. *tritici* (Hu et al., 2018). The larger number of effector transcripts at the later time point is likely due to an increased presence of *E. pisi* cells in the samples, although R and PR accessions were more successful in suppressing effector expression over time. Many of these predicted effectors showed protein structures similar to proteins found in other pathogenic fungi such as *Puccinia graminis, Magnaporthe oryzae*, *Rhizoctonia solani, Phytophthora infestans*, and *Blumeria graminis*.

Most transcriptomic studies on plant host responses to pathogens focus primarily on complete resistance (described as having an incompatible reaction with the pathogen) and complete susceptible accessions, often overlooking the nuances between complete and partial resistance or susceptibility - key factors for developing more durable plant resistance to pathogens. In this study, we selected four *L. sativus* accessions (R, PR, PS, and S) from a worldwide collection previously phenotyped for response against *E. pisi* using detached leaves (Martins et al., 2023). These phenotypes were confirmed in the present study, using whole seedlings, with a progressive increase in macroscopic DS from the R, PR, PS, to the S accession, ranging from 0 to 35% at 7 dai and from 1.5 to 77.5% at 14 dai. Although the R accession displayed a compatible reaction with *E. pisi* at 14 dai (IT=3, DS=1.5), our transcriptomic analysis focused on the period from 0 to 72 hai - a period where no visible macroscopic differences were observed among accessions. During these early infection phases, the four accessions showed different approaches in the infection response. *L. sativus* is known for its remarkable resilience to pests and diseases, offering valuable diverse genetic resistance against major fungal diseases in legumes (Vaz Patto et al., 2006; Vaz Patto et al., 2014; Almeida et al., 2015). Indeed, even the most susceptible *L. sativus* accession in our study showed lower DS compared to the pea cultivar ‘Messire’, used as a susceptible control, highlighting *L. sativus* generally higher resistance to *E. pisi* compared to pea.

### 4.1. Different *Lathyrus sativus* accessions activate common mechanisms in response to *Erysiphe pisi* inoculation

We observed common defence responses against *E. pisi* infection across PS, PR and R accessions, with enrichment in stress and/or stimulus response processes in all time points after inoculation. This indicates that accessions with contrasting resistance phenotypes activate common mechanisms in response to pathogen-induced stress, though the intensity (FC) and timing (constitutively, early, or late during infection) of these responses differ. The key common resistance mechanisms in *L. sativus* against *E. pisi* involve antifungal proteins, cell wall reinforcement, ROS-mediated defence, and SAR. At early infection stages (12 hai), starch and glucan biosynthesis processes were downregulated across PS, PR and R accessions, indicating an early metabolic shift from growth-related processes to defence responses under pathogen attack (Chaliha et al., 2018), as also reported in *M. truncatula* infected by *E. pisi* (Gupta et al., 2020).

Examples of defence-related genes common to PS, PR and R accessions included Bowman-Birk type proteinase inhibitors (BBIs), peptidyl-prolyl cis-trans isomerase *FKBP62*, BURP domain proteins (*RD22*), polygalacturonase inhibitors (PGIPs), mannitol dehydrogenases, and flavin-containing monooxygenases. BBIs are well-known for their antifungal properties, blocking proteases secreted by pathogens, preventing the degradation of plant proteins and hindering pathogen growth (Gitlin-Domagalska et al., 2020). BBIs represent a well-conserved defence strategy common in legumes, which seems to be crucial for resisting *E. pisi* infection (Gitlin-Domagalska et al., 2020).

Peptidyl-prolyl cis-trans isomerases FKBP62 were also strongly upregulated in R, PR, and PS compared to S accession. This protein participates in several defence mechanisms, including the accumulation of callose in the cell wall, which reinforces structural barriers against pathogen invasion (Pogorelko et al., 2014; Ge et al., 2022). This mechanism resembles the *er1* (*PsMLO1*) gene functions, where the loss of function of the MLO1 protein in pea leads to increased protein cross-linking in host cell walls, creating a physical barrier that prevents penetration by the pathogen (Iglesias-García et al., 2015). *MLO* transcripts were not differentially expressed in this study, consistent with findings from other transcriptomic analyses of legume-*E. pisi* interactions (Gupta et al., 2020; Bhosle & Makandar, 2021).

BURP domain proteins (RD22), typically associated with abiotic stress responses, may also play a role in biotic stress through cell wall reinforcement (Yu et al., 2022). Similarly, polygalacturonase-inhibiting proteins (PGIPs) enhance structural defences by inhibiting fungal polygalacturonases, enzymes that degrade cell wall pectin to allow pathogen entry (Kalunke et al., 2015). Indeed, we detected effectors specialised in cell wall hydrolysis such as an alpha-glucosidase (DN9453) expressed in all samples, a hemicellulose-degrading Mannan endo-1,6-alpha mannosidase DCW1 (DN4717) expressed in PR and S accessions (Lee et al., 2023), a pectin lyase-like protein (DN174), and a secreted glycosidase (DN9922) both expressed in all accessions but the PR accession. By blocking these enzymes, PGIPs prevent cell wall breakdown, fortifying the plant’s structural defences against *E. pisi*. In legumes, PGIPs are involved in interactions with apoplastic peroxidases, leading to increased lignin production (WANG et al., 2012). In *L. sativus*, these proteins were upregulated in all accessions compared to the S, with higher expression in the R and PR accessions.

Mannitol dehydrogenases were highly upregulated across all time points in PS, PR and R accessions in response to *E. pisi* infection. Mannitol, a polyol commonly found in fungal spores, fruiting bodies, and mycelia, is secreted by fungi (including Erysiphales) to neutralize reactive oxygen species (ROS), which mediate plant host defences (Solomon et al., 2007; Meena et al., 2015). To counter this, pathogen-induced plant mannitol dehydrogenases break down fungal mannitol, restoring ROS activity and preventing pathogen spread (Meena et al., 2015). Notably, we observed elevated expression of mannitol dehydrogenases in the host R accession at 48 and 72 hai, highlighting their role in enhancing ROS-mediated defences. Accordingly, at 72 hai, several biological processes involved in oxidative stress responses were upregulated across all accessions compared to the S accession.

Flavin-containing monooxygenase 1 (*FMO1*), another key gene upregulated across all comparisons, is a known marker of SAR, a long-lasting defence mechanism that is triggered by pathogen-induced cell death and protects the plant from several pathogens beyond the immediate fungal threat (Durrant & Dong, 2004; Mishina & Zeier, 2006; Olszak et al., 2006).

When analysing the expression patterns of predicted NLRs, *RUN1*, *RPP13-like protein 1*, *RGA2* and *NRG1.1* were particularly highly expressed across all *L. sativus* accessions. Their role in defence against various powdery mildew pathogens is well-documented (Cheng et al., 2018; Marimon et al., 2020; Bhosle & Makandar, 2021; Massonnet et al., 2022; Wang et al., 2014; Zhang et al., 2022), highlighting their potential involvement in *L. sativus*’ response to *E. pisi.* As example, *NRG1.1*, also known as *NRG1A*, is an extensively studied helper CC_R_-NLR characterised by its resistance to powdery mildew 8 (RPW8) domain, which controls resistance to a broad range of powdery mildew pathogens (Xiao, 2004). *NRG1* is required for signal transduction of many sensor TIR-NLRs such as *RUN1*, and can cause autoimmunity when mutated, highlighting its importance in triggering HR (Xiao, 2004; Wu et al., 2019; Sun et al., 2021). However, a helper NLR requires a sensor NLR to recognise its specific effector first to promote cell death (Sun et al., 2021), thus its heightened expression should not be vital to define resistance or susceptibility to a pathogen. Another example of an NLR that maintained high expression in all samples was an *RPS2-like* CC_G10_-NLR (g20538.t1), which was previously found in linkage disequilibrium with a SNP marker associated with *L. sativus* resistance to *E. pisi* (Martins et al., 2023).

### 4.2. R and PR accessions prioritised physicochemical barriers, secretion of antifungal compounds, and immune signal pathways

Transcriptome reprogramming in response to *E. pisi* infection revealed many specific similarities between R and PR accessions, suggesting that both accessions converge on common gene expression patterns to combat *E. pisi* infection. The common defence mechanisms in R and PR accessions involve a multifaceted strategy including physical barriers, chemical defences, antifungal proteins, ROS quenching, NLR expression, HR, and hormone signalling pathways, with the R accession showing a faster and stronger activation than the PR accession.

The high expression observed for genes related to processes like cell wall reinforcement, phenylpropanoid metabolism, and flavonoid biosynthesis both in R and PR accessions, has also been reported in *L. cicera*, *P. sativum* and *M. truncatula* responses to *E. pisi* (Bhosle & Makandar, 2021; Foster-Hartnett et al., 2007; Santos et al., 2020). One of the key enzymes in the phenylpropanoid pathway, 4-coumarate-CoA ligase (CCL1), was already upregulated before inoculation (at 0 hai), increasing its expression at 12, and 72 hai, in both PR and R accessions. *CCL1* is crucial for the biosynthesis of lignin and flavonoids (Cao et al., 2020; Li et al., 2020), which play important roles in plant defence. Lignin strengthens the plant cell wall and acts as a physical barrier to pathogen penetration (Lee et al., 2019; Saberi Riseh et al., 2024), while flavonoids provide chemical defence through antifungal activity, ROS quenching, and triggering HR (Mierziak et al., 2014). Flavonoids can also inhibit pathogen enzymes by chelating metals necessary for their activity (Mierziak et al., 2014). Therefore, the elevated expression of *CCL1* in PR and R accessions likely contributes to both chemical and physical defences against *E. pisi* infection.

The expression of genes involved in trichome morphogenesis and plant epidermis development such as *Protein CPR-5* and *Protein SCAR2* may play a role in the R and PR accessions resistance to *E. pisi* both constitutively (0 hai) and in the early stages of infection (12 hai). *CPR-5* controls trichome cell cycle transition and activates plant effector-triggered cell death (Peng et al., 2020). *SCAR2* is required for epidermal morphogenesis and regulates trichome branch (Basu et al., 2005; Zhang et al., 2005). Trichomes act as dynamic passive barriers and prevent spores and other microbial elements from reaching the leaf surface (Karabourniotis et al., 2020). Beyond their structural role, trichomes contain phenolic compounds, such as flavonoids, which further enhance the plant’s defence mechanisms. Studies in Cucurbitaceae have shown that increased trichome density and polyphenol accumulation in the epidermis are associated with reduced susceptibility to the stem blight (*Didymella bryoniae*) (Rennberger et al., 2017). Similarly, resistance to rust (*Puccinia helianthi*) in sunflower is associated with coumarin and other phenolic compounds’ accumulation on leaf surface that impair germ tube growth (Prats et al., 2007a).

In addition to physical or physicochemical barriers, both R and PR accessions showed high expression of genes encoding antifungal proteins like Kunitz-type trypsin inhibitors (KTIs), which are predicted secreted proteins with antifungal activity (Huang et al., 2010; Cai et al., 2018). In *L. cicera*, a KTI was identified as a candidate gene for resistance against *E. pisi* (Santos et al., 2020), supporting their importance in *Lathyrus* species’ defence strategies.

A shared set of DEGs involved in plant immunity was upregulated in both R and PR accessions. An interesting example is the receptor-like protein Cf-9. This receptor-like protein (RLP) has extracytoplasmic leucine-rich repeats eLRRs that confer disease resistance through recognition of fungal effectors (ETI), resulting in the activation of HR (van der Hoorn et al., 2005; Jamieson et al., 2018). Therefore, though structurally similar to PRRs, RLP Cf-9 behaves as a NLR (Jamieson et al., 2018). RLP Cf-9 was highly expressed in R and PR accessions, especially at 0 and 12 hai, suggesting a fast *E. pisi* recognition, immune signalling and prevention of pathogen progression, in these accessions. Although HR has not been observed macroscopically in the studied *L. sativus* accessions infected by *E. pisi*, HR is an effective mechanism against biotrophic pathogens, including legume powdery mildew, as observed macro- and microscopically for *Pisum* species (Fondevilla et al., 2006, 2007; Fondevilla & Rubiales, 2012) and *M. truncatula* (Prats et al., 2007b). Therefore, detailed histological studies are still needed to investigate the role of HR in the *L. sativus* response against *E. pisi*.

Regarding NLR genes potentially involved in the ETI response, *RUN1* (g15597.t1) and *RGA1* (g11157.t1) showed significantly higher expression levels in the R and PR accessions. Notably, *RGA1* has also been reported to be upregulated in other plant-aerial pathogen interactions, such as *Oryza sativa* infected by *Rhizoctonia solani* (sheath blight) and *Triticum aestivum* infected by *Puccinia striiformis* f. sp. *tritici* (rust) (Zheng et al., 2020; Durgadevi et al., 2021).

Both R and PR accessions upregulated genes involved in the JA signalling pathway, a stress-responsive hormone produced during pathogen attacks that activate key defence mechanisms (J. Yang et al., 2019). For example, the allene oxide cyclase (AOC), a key enzyme in the biosynthesis of JA, was upregulated in both accessions at all time points. Notably, recent studies on *Medicago* spp. identified *AOC* as pathogen-responsive (Yang et al., 2023).

At 48 hai, R and PR accessions also shared upregulated genes previously linked to abiotic stress responses, particularly water-related stress, which persisted into 72 hai. An example was the *NAC domain-containing protein JA2L*, that was minimally expressed in PS accession but exhibited significantly higher expression in R and PR accessions at all time points, including at 6 and 24 hai. In particular, the R accession displayed a steady increase in *JA2L* expression from 0 to 72 hai. In tomato, *JA2L* targets genes involved in SA metabolism, a key component of plant defence responses (Kotera et al., 2023). Interestingly, in the R accession, the regulation of stomatal movement was significantly upregulated compared to S, particularly at 0, 12, and 48 hai, stressing its importance in R’s defence strategy. In parallel, SA biosynthesis in R and PR accessions was reinforced through the upregulation of benzyl alcohol O-benzoyltransferases, which in conjugation with peroxisomal β-oxidative pathway contribute to pathogen signal-induced SA production (Kotera et al., 2023). In our study, those genes were highly upregulated at 12, 48, and 72 hai in R and PR, highlighting their role in strengthening immune responses against pathogen invasion.

### 4.3. The R accession showed a robust constitutive physicochemical defence response

In addition to the common genes and defence mechanisms between R and PR accessions, each of these accessions displayed unique molecular responses to *E. pisi*, contributing to their varying DS values. While physical and chemical barrier reinforcement was common between the accessions, variations in the type, number of DEGs, timing, and intensity (FC) distinguished their response. In the R accession, it was clear that the defence strategy combines early and rapid reinforcement of structural barriers with sustained chemical defences and stress responses. Before inoculation, upregulated BPs uniquely found in the R accession were related to lignin biosynthesis and the phenylpropanoid pathway, reinforcing its structural barriers against possible pathogen attack. Additionally, the *hypersensitive-induced response protein 1* gene involved in HR activation (Zhou et al., 2010) was only upregulated in the R accession, not only prior to inoculation, but also across all infected time points, suggesting a role of HR in R defence. Moreover, the CC_G10_-NLR *SUMM2* (g31324.t1) was upregulated at 0, 12 and 48 hai in R accession. *SUMM2* does not directly sense the pathogen effectors; instead, it monitors the phosphorylation status of the plant calmodulin-binding receptor-like cytoplasmic kinase 3 and can trigger autoimmunity in specific knockout backgrounds (Zhang et al., 2017). This potent autoimmune ability to induce HR on its own is characteristic of CC_G10_-NLRs, also known as the autonomous NLR clade (Lee et al., 2021).

Antifungal proteins such as *eugenol synthase 1* and *thaumatin-like protein 1* appear to play a role, particularly in the R accession response to *E. pisi*. Eugenol synthase is an enzyme responsible for the biosynthesis of eugenol, a volatile with antifungal activity (Anand et al., 2016; Ulanowska & Olas, 2021). Expression analysis of *eugenol synthase 1* revealed that this gene is exclusively expressed in the R accession at 6 hai, while at all other analysed time points, it is expressed in the R, PR, and S accessions, albeit at lower levels in the S accession. Thaumatin-like proteins (TLPs) are part of the pathogenesis-related protein family and play a crucial role in providing resistance against various fungi, including *E. pisi* and other pathogens that infect legumes (Jayaprakash et al., 2021; Zhou et al., 2023; Feng et al., 2024). Here, TLP1 was upregulated in R accession only at 12 hai.

Upon *E. pisi* infection at 12 hai, alongside the continued emphasis on physical and chemical defences, the R accession uniquely showed high expression of genes related to epigenetic regulation, such as DNA replication, conformation, and unwinding. This suggests that DNA repair, transcription activation, and chromatin reorganization may be involved in the plant’s defence response to the pathogen. In the R accession at 12 and 72 hai, DNA unwinding mediated by DNA replication licensing factors were more expressed, particularly *mini-chromosome maintenance* (*MCM*) genes. MCM proteins ensure that genomic DNA is replicated completely and accurately during the S phase of the cell cycle (Tuteja et al., 2011). Since pathogen exposure affects plant growth, it may directly or indirectly affect cell cycle regulation by altering the endoreduplication process. This may result in DNA replication perturbation and cell death (Tuteja et al., 2011). In pea, the PsMCM6 was suggested to function as a helicase, aiding in unwinding the secondary structures of mRNA in stress-responsive genes (Tuteja et al., 2011). In Arabidopsis, *MCM7* has been reported to be expressed during root-knot and cyst nematode infections (Huang et al., 2003). Moreover, previous studies have shown that the activation of PTI, ETI, and SAR depends on epigenetic regulation of gene expression (Chen et al., 2017). Thus, these mechanisms are likely important for a robust and systemic response to *E. pisi* in the R accession.

At later stages of infection (72 hai), the R accession transitioned to sustaining resistance through mechanisms that include oxidative stress management, osmotic stress responses, and abscisic acid (ABA) signalling. At this time point, genes involved in heat and oxidative stress responses, such as *peroxidase 4*, and *heat shock proteins,* were strongly upregulated in R accession. Additionally, protein folding, and detoxification mechanisms became more prominent, indicating that cellular homeostasis under stress is essential for adaptation and survival. Peroxidases, as members of the pathogenesis-related protein family, play a crucial role in maintaining redox homeostasis within plant cells (Sellami et al., 2022; dos Santos & Franco, 2023). Besides their role in cell signalling after infection, peroxidases contribute to plant defence by polymerizing macromolecules that, once deposited on the extracellular surface, promote cell wall strengthening (dos Santos & Franco, 2023). Additionally, peroxidases can catalyse the oxidative degradation of phenolic compounds in the damaged cell regions caused by pathogens (dos Santos & Franco, 2023). *Peroxidase 52* was reported as showing very high transcript expression in resistant pea genotypes against *E. pisi*, highlighting the potential role of peroxidases in enhancing resistance mechanisms against this pathogen.

Many different heat shock proteins (HSPs) and related transcription factors were uniquely and highly upregulated in the R compared to S, particularly at 72 hai. Initially discovered in response to temperature increases, HSPs play diverse roles in plants by acting as molecular chaperones, facilitating protein assembly, stabilisation, and maturation (ul Haq et al., 2019). Additionally, HSPs enhance membrane stability and help detoxify ROS by positively regulating antioxidant enzyme systems (ul Haq et al., 2019). As a result, they are crucial in plant responses to various abiotic and biotic stresses, including pathogen infections (Park & Seo, 2015; ul Haq et al., 2019). The role of HSPs in the resistance to *E. pisi* was also reported in *L. cicera* and *P. sativum* (Curto et al., 2006; Santos et al., 2020; Bhosle & Makandar, 2021). Interestingly, *E. pisi* also expressed HSPs Rot1 (DN4492) and chaperone J-domain-containing protein (DN8317) to protect itself from plant defence responses.

### 4.4. The PR accession responds to *Erysiphe pisi* by activating biotic stress-specific processes from the earliest infection stage

PR-specific molecular responses against *E. pisi* mainly relied on key BPs related to biotic stress defence and signalling, including the expression of NLR genes. Before inoculation, the PR accession exhibited a unique defence-related transcriptome, distinct from both R and PS accessions compared to the S. Prominent defence-related genes constitutively upregulated in PR include 15 NLRs, suggesting this accession is primed for an effective response even before pathogen exposure.

At 12 hai, the PR accession continued to exhibit the strongest defence-focused response, marked by increased activity in processes related to biotic stimuli, including fungal infection. This robust response, sustained from 0 to 12 hai, highlights the PR accession’s ability to react quickly to pathogen presence. However, as the infection progresses to 48 and 72 hai, the transcriptomic activity in defence-related processes declines. Thus, the overall defence response in the PR accession appeared to be front-loaded, reducing intensity as the infection progressed, which might limit its long-term effectiveness against sustained pathogen pressure, phenotypically distinguishing this accession from R.

Contrary to the previous paradigm, NLRs have recently shown not to always promote complete resistance, and can instead be agents of partial resistance such as non-race-specific resistance like *Pik*, a CC-NLR conferring complete or partial resistance to *Magnaporthe oryzae* in rice (Varden et al., 2019). Other examples include the *I2*, a CC-NLR conferring resistance to *Fusarium oxysporum* f. sp. *lycopersici* in tomato, or when mutated in specific residues, conferring partial resistance to *Phytophthora infestans* (Giannakopoulou et al., 2015), and the *RGA5* NLR which confers partial resistance to *Blumeria graminis* in wheat (Liu et al., 2024). Additionally, there is another way NLR proteins contribute towards partial resistance: by mis-regulation of NLR gene expression; meaning enough NLR activation needs to occur for a complete resistance response (Fick et al., 2022b). Therefore, if there is not sufficient NLR expression, there is poor detection of pathogen presence, leading to lower levels of immune response activation, and thus the plant can acquire a partial resistance phenotype (Fick et al., 2022a).

### 4.5. PS accession delayed and generalised defence responses to *Erysiphe pisi*

PS accession exhibited a constitutive response pattern mainly focused on general environmental stress rather than on pathogen-specific defence, reflecting a potentially weaker baseline defence system when compared to the PR and R accessions. Defence-related processes and responses to external biotic interactions were more evident at 48 hai, occurring later than in the R and PR accessions. The NLR *At4g27190* (g14462.t1) was upregulated in the PS samples at 0, 48, and 72 hai. Notably, four homologs of *At4g27190* were found to be upregulated in a resistant *Gerbera hybrida* accession compared to a susceptible accession when challenged with the powdery mildew causal agent *Podosphaera xanthii* (syn. *Sphaerotheca fusca*) (Bhattarai et al., 2020).

### 4.6. The *Erysiphe pisi de novo* transcriptome helped effector prediction

Effectors are crucial for powdery mildew virulence, as they interact with host defence-related proteins to weaken host resistance and promote successful fungal colonisation (Hu et al., 2018). Despite focusing only on 40 effectors, we identified 297 potential *E. pisi* effectors, a number that falls between the 681 effectors reported by Bhosle & Makandar (2021) for isolate Ep01, and the 167 effectors identified by Sharma et al. (2019) for isolate Palampur-1, both infecting *P. sativum*. Notably, 27 of the 167 effectors in Sharma et al.’s study correspond to 23 of our identified effectors.

From the 40 *E. pisi* effectors with Foldseek hits selected in the present study, 16 were expressed in all accessions in at least one inoculated time point. One of them was a 7-dehydrocholesterol reductase (DN10505), an enzyme known to play a significant role in the sterol biosynthesis pathway, which is involved in *Phytophthora capsici* development and zoospore virulence (Wang et al., 2022). Another example is the SnodProt1 (DN1473), a member of the cerato-platanin protein family that is required for the virulence of different pathogens (Jeong et al., 2007; Zhang et al., 2017a; Nasir et al., 2018). This protein can also function as a plant defence elicitor (Zhang et al., 2017a; Nasir et al., 2018). We also detected two other secreted effector proteins (DN1391, DN144) in all accessions, both containing a ribonuclease/ribotoxin domain. Fungal ribotoxins were observed in other *E. pisi* studies (Gupta et al., 2020; Sharma et al., 2019) and typically induce programmed cell death. Additionally, these ribotoxins can act as elicitors, triggering HR depending on plant genotype (Yin et al., 2022).

Among the 40 studied *E. pisi* candidate effectors, we found six specific effectors only expressing in S and PS. Among them, we identified a fungal-specific transcription factor (DN8238) at 72 hai which could be modulating the expression of relevant pathogenicity-related or even host resistance genes; a secreted serine-threonine phosphatase (DN708) at 48 and 72 hai that can alter the activation state of defence host proteins; a V-type proton ATPase (DN5816) at 48 and 72 hai to provide chemical energy for host-pathogen interactions; an histone acetyltransferase ELP3 (DN4049) in all time points for epigenomic gene modulation, and a pre-rRNA processing protein (DN1179) at 48 and 72 hai denoting high translation activity. These effectors seem to contribute towards a broad transcriptional and translational activity but do not point towards any specific pathogenicity strategy. Low specificity is frequently a characteristic of the most abundant effectors, as targeting multiple host targets enhances the likelihood of blanketing the entire plant defence-signalling network, thereby promoting disease (Khan et al., 2018).

The *E. pisi* genome and its effectorome in interaction with *L. sativus* remained largely unexplored. Furthermore, effectors are an ever-changing family, with pronounced differences even among strains (Khan et al., 2018). Thus, to explore *E. pisi*’s effectorome, we generated a *de novo E. pisi* transcriptome containing 20,608 transcripts from 8,031 genes. This number of genes is within the expected range for powdery mildew fungi, which usually have a smaller genome than other saprotroph fungi, usually between 6,046 to 8,470 genes due to biotrophism (Zaccaron et al., 2023). On the host side, this is the first transcriptomic study using the recently released high-quality *L. sativus* genome (Vigouroux et al., 2024). The availability of this genome provided a robust reference for mapping transcriptomic data, enabling precise identification and annotation of DEGs. Despite the high overall functional annotation rate, 24% of DEGs were classified as ‘no annotation’, suggesting that these may represent *L. sativus*-specific genes, possibly involved in unique defence mechanisms against *E. pisi* that have yet to be described.

## 5. Conclusion

This study provides valuable insights into the complex molecular interactions and defence mechanisms in *L. sativus* against *E. pisi*, underscoring the genetic diversity in pathogen responses across accessions with contrasting resistance levels. By identifying genes and pathways involved in different resistance mechanisms, breeders can employ pyramiding techniques to combine and integrate them into a single genotype to create varieties with broader resistance spectra. Moving forward, future research should focus on the functional validation of the most promising candidate genes, including NLRs, and the assessment of effector functions. We identified NLRs consistently more expressed in resistant accessions, which could be sensing, directly or indirectly, *E. pisi* effectors. NLRs g11147.t1 (*RGA3*), g14573.t1 (*RPP13-like protein 1*), g18217.t1 and g18222.t1 (*RPM1*), and g29548.t1 (*RUN1*) could contribute to *E. pisi* resistance, and we recommend their functional validation in future studies. The characterisation of *E. pisi* effectors and their interactions with *L. sativus* defence genes will be crucial for understanding the molecular basis of powdery mildew disease and developing effective resistance breeding strategies and tools, such as marker-assisted selection. The insights gained from *L. sativus* can also be utilised to enhance resistance in other economically important legumes susceptible to *E. pisi*, such as pea. By improving disease resistance in these crops, we can boost legume yields while promoting more sustainable agricultural practices.

## Acknowledgements

We thank the U.S. Department of Agriculture for providing the *Lathyrus sativus* PI accessions. We acknowledge Manuel Alejandro Vaquero (IAS-CSIC, Spain) for assisting with maintenance and multiplication of *Erysiphe pisi* isolates, and María José Cobos (IAS-CSIC, Spain) for technical support. We also acknowledge Joe Win (TSL, UK) for advising on effector prediction.

## Funding

This research was funded by Fundação para a Ciência e Tecnologia (FCT, Portugal) through the research project 2022.08266.PTDC R&D (DOI: 10.54499/2022.08266.PTDC), Research Unit GREEN-IT— Bioresources for Sustainability (UIDB/04551/2020 DOI: 10.54499/UIDB/04551/2020 and UIDP/04551/2020 DOI: 10.54499/UIDP/04551/2020) and the LS4FUTURE Associated Laboratory (LA/P/0087/2020). Spanish AEI project PID2023-146215OB-I00 is also acknowledged. RMM was supported by the FCT grant UI/BD/151214/2021, CS received support through the 2017.00198.CEECIND (DOI 10.54499/CEECIND/00198/2017/CP1428/CT0002) research contract and STL was supported by the research contract 2022.00163.CEECIND (DOI 10.54499/2022.00163.CEECIND/CP1725/CT0021).

## Data availability

The RNA-seq data and corresponding metadata is available at the National Center for Biotechnology Information <u>BioStudies - EMBL-EBI</u>) under accession number E-MTAB-14647. Any additional data will be made available on request.

## Declaration of competing interest

The authors declare that they have no known competing financial interests or personal relationships that could have appeared to influence the work reported in this paper.

## Notes

### Competing Interest Statement

The authors have declared no competing interest.

