## Supplementary material for "Comparative transcriptomics of *Lathyrus sativus* reveals accession-specific resistance responses against *Erysiphe pisi*": Fig. S

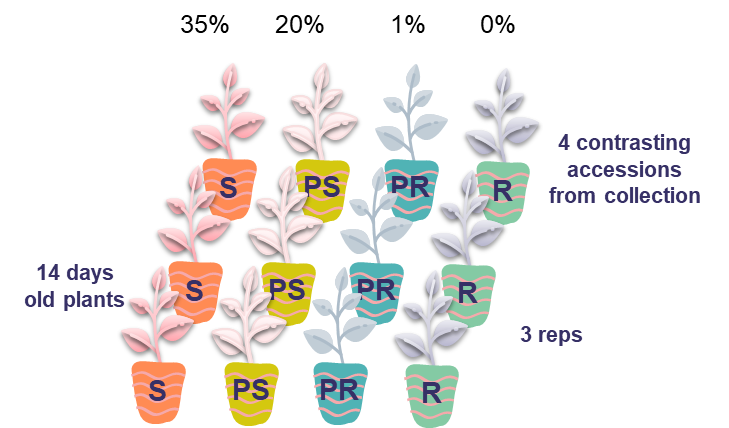

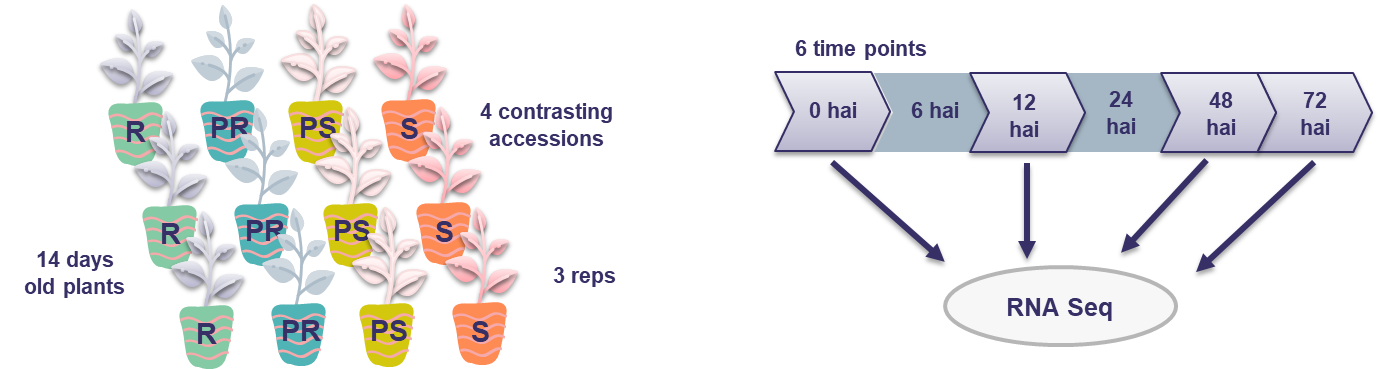
**Supplementary Figure 1** Experimental design of the RNA Seq experiment. Three biological replicates of four contrasting *Lathyrus sativus* accessions against *Erysiphe pisi* (S – PI426890, PS – PI426882, PR – PI221467_A, R - PI268478) were inoculated. Infected leaf material from 14 days-old plants was collected at 0, 6, 12, 24, 48, and 72 hai, and the 0, 12, 48, and 72 hai time points were selected for RNA Seq.

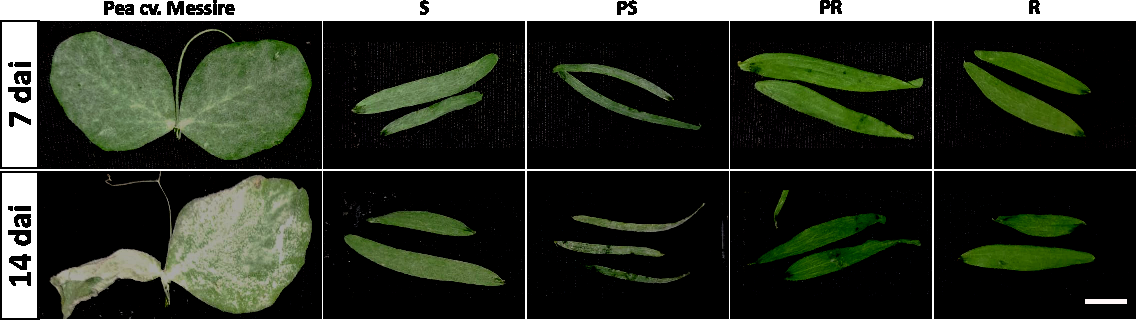

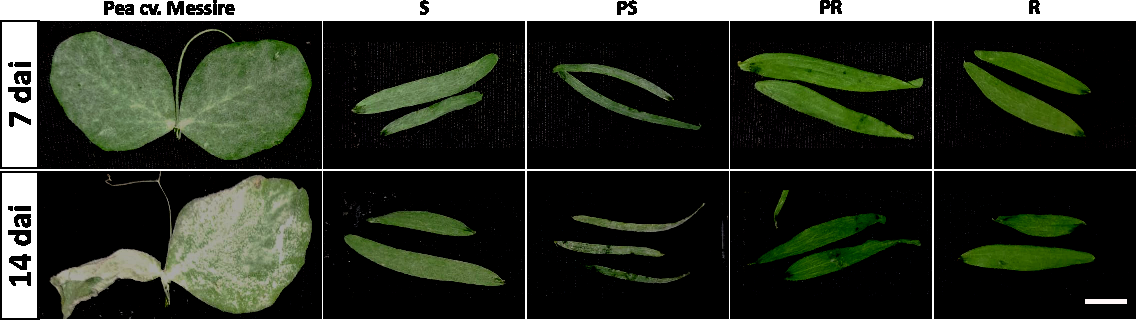

88%

35%

22.5%

0.5%

0%

95%

77.5%

72.5%

3%

1.5%

**Supplementary Figure 2** Disease symptoms of the *Lathyrus sativus* accessions S (Susceptible - PI426890), PS (Partial Susceptible - PI426882), PR (Partial Resistant - PI221467_A), and R (Resistant - PI268478) at 7 and 14 days after inoculation (dai) with *Erysiphe pisi*. The pea cv. ‘Messire’ was used as a susceptible control for inoculation. The numbers depicted on the bottom left corner represent the average disease severity (DS) across replicates. Scale: 1 cm.

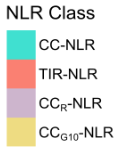

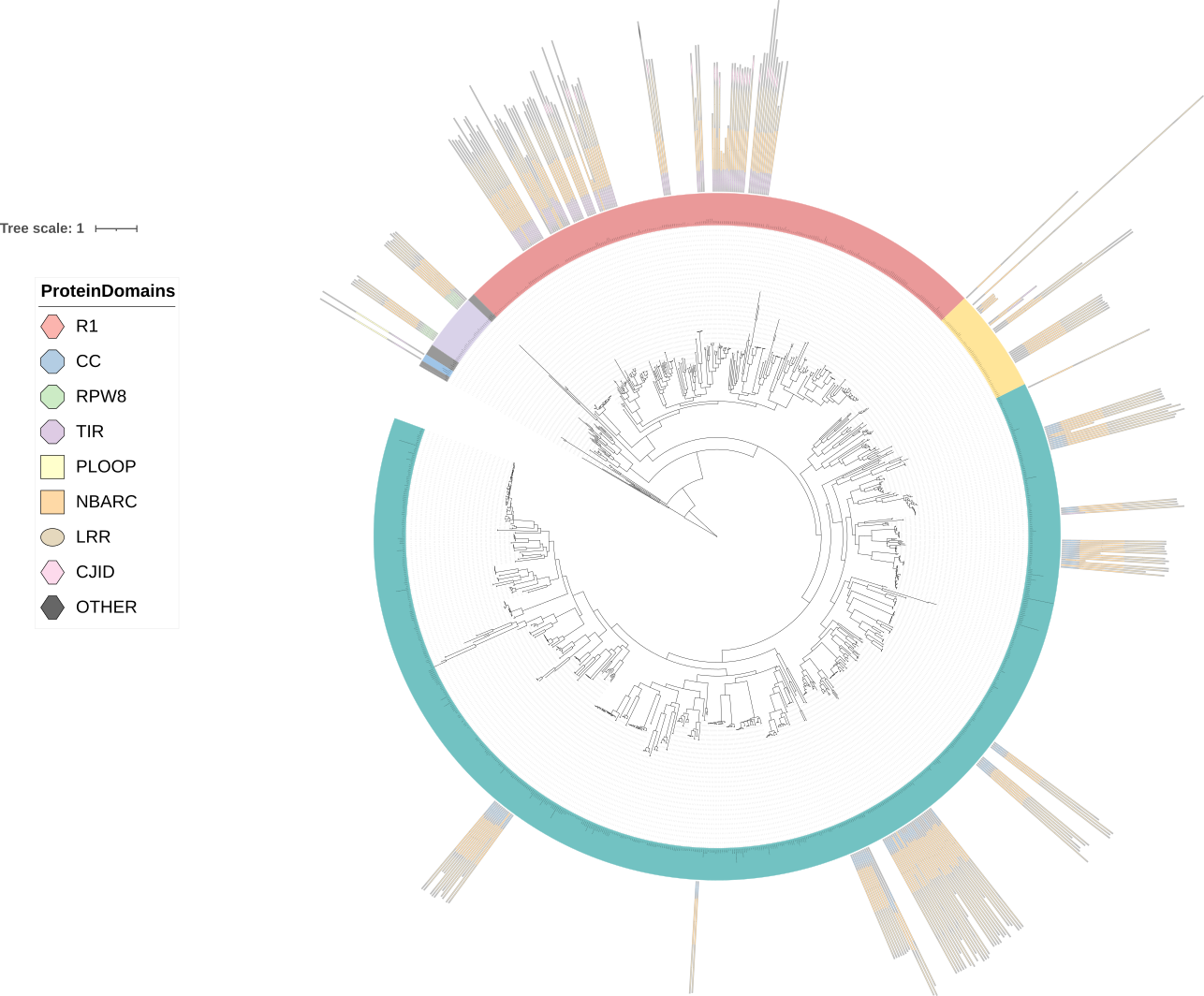
**Supplementary Figure 3** *Lathyrus sativus* NLR phylogenetic tree including NLRs from the RefPlantNLR dataset. This image was obtained using the iTOL website. The legend depicts common NLR domains predicted by NLRtracker. NLR classes are represented with the following colour code: CC-NLR – turquoise; TIR-NLR – salmon; CC_G10_-NLR – yellow; CC_R_-NLR – lilac.

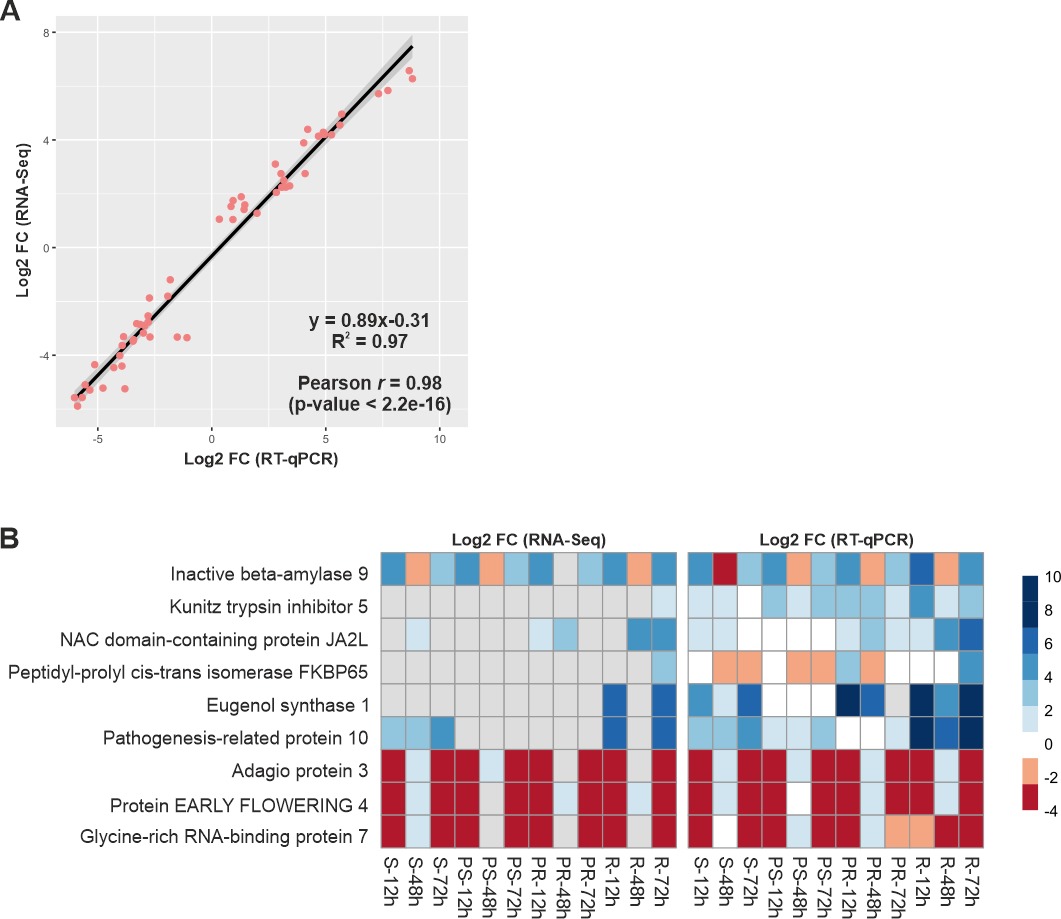

**Supplementary Figure 4** Comparisons of target gene expression by RT-qPCR for technical validation. (A). Linear regression and Pearson correlation analysis between RT-qPCR and RNA-seq results (*r* = 0.933) for the selected genes. X-axis numbers represent the fold change values of RT-qPCR results. Y-axis numbers represent the fold change values of RNA-seq results. (B) Heatmap representing log_2_ fold change expression values from RNA-seq and RT-qPCR for the selected genes.
